# A statistical learning method for simultaneous copy number estimation and subclone clustering with single cell sequencing data

**DOI:** 10.1101/2023.04.18.537346

**Authors:** Fei Qin, Guoshuai Cai, Feifei Xiao

## Abstract

The availability of single cell sequencing (SCS) enables us to assess intra-tumor heterogeneity and identify cellular subclones without the confounding effect of mixed cells. Copy number aberrations (CNAs) have been commonly used to identify subclones in SCS data using various clustering methods, since cells comprising a subpopulation are found to share genetic profile. However, currently available methods may generate spurious results (e.g., falsely identified CNAs) in the procedure of CNA detection, hence diminishing the accuracy of subclone identification from a large complex cell population. In this study, we developed a CNA detection method based on a fused lasso model, referred to as FLCNA, which can simultaneously identify subclones in single cell DNA sequencing (scDNA-seq) data. Spike-in simulations were conducted to evaluate the clustering and CNA detection performance of FLCNA benchmarking to existing copy number estimation methods (SCOPE, HMMcopy) in combination with the existing and commonly used clustering methods. Interestingly, application of FLCNA to a real scDNA-seq dataset of breast cancer revealed remarkably different genomic variation patterns in neoadjuvant chemotherapy treated samples and pre-treated samples. We show that FLCNA is a practical and powerful method in subclone identification and CNA detection with scDNA-seq data.

## INTRODUCTION

In cancer, a small population of cancer stem cells evolves into a malignant mass of tumor cells, which then diverges and forms distinct subclones, contributing to intra-tumor heterogeneity (ITH). The level of ITH is associated with tumor progression and is sensitive to clinical treatments (1). Intrinsic mechanistic processes, such as inherent genomic variation, clonal competition, and tumor-host interactions, contribute to ITH (2–5). Therefore, accurate assessment of ITH and identification of subclones is essential to understand the mechanisms of tumor progression and resistance to therapy (6).

Most existing studies (7–11) characterize clonal diversity using bulk DNA sequencing, which was limited in only reporting an average signal from a complex population of cells (12). The emerging single cell sequencing (SCS) technology enables the assessment of ITH on a single-cell basis (13, 14), providing single cell DNA or RNA sequencing (scDNA/RNA-seq) information to reveal cellular evolutionary relationship (15). Subclonal populations can be identified from the same tumor tissue using SCS data, allowing for the inference of tumor evolution and providing insights into the development of targeted therapy for different tumor types (16). We take breast cancer for illustration, which is the most common malignancy in adult women. Triple-negative breast cancer (TNBC) is an important class of breast cancer which constitutes 12-18% of breast cancer patients (17), and many studies have shown that patients with TNBC harbor high levels of somatic mutations (14, 18), which partially result in extensive ITH. Neoadjuvant chemotherapy (NAC) is the standard therapy for TNBC patients who show low level of the estrogen, progesterone and HER2 receptors and are not eligible for hormone or HER2-targeted therapy (17). Still, due to chemoresistance, poor overall survival performance is observed in about 50% TNBC patients (17). Identifying subclones and detecting mutations in this subpopulation is critical to untangle the molecular mechanisms of chemoresistance in these patients.

Cells in a subpopulation share genetic variant characteristics (19), with chromosomal copy number aberration (CNA) being one of the most important genetic variants, which is the gain or loss of DNA segments. CNAs stimulate the stemness of other tumor cells to new cancer stem cells, ultimately giving rise to clonal evolution in cancers (20). Therefore, CNAs serve as good biomarkers to assess ITH and identify subclones. Basically, the detection of CNAs is to find the breakpoints or boundaries of copy number regions from chromosomal segments. Though great efforts have been made in CNA detection methodology for scDNA-seq data, it is challenging to detect CNAs in such data due to shallow and uneven depth of coverage (21). We use the existing methods applied in our study for brief illustration. SCOPE (22) was recently developed for copy number estimation with whole genome scDNA-seq data which uses a Poisson latent factor model for normalization and an expectation-maximization (EM) algorithm to estimate copy number profile. HMMcopy was designed for array comparative genomic hybridization (aCGH) and next generation sequencing (NGS) data (23, 24) using a Hidden Markov Model, and has also been extensively applied in SCS data (25). Relevant to our proposed method in this study, Rojas & Wahlberg (26) used a fused lasso method and converted the problem into a convex optimization problem to detect chromosomal breakpoints. However, studies based on these methods used an independent statistical model to detect copy number profile in the first step, followed by classical clustering methods (e.g., Hierarchical (27)) for subclone identification in downstream analyses. This two-step framework will generate spurious results due to the carry-over of noisy signals in the first copy number profiling step into the process of subclone clustering. We herein developed a method to achieve accurate copy number profiling and subclone clustering without copy number state calling as a middle step. It was the first time that the fused lasso model was explored in copy number profile based subclone clustering.

We herein developed FLCNA, a CNA detection method based on the fused lasso model, which simultaneously identifies subclones with scDNA-seq data. The FLCNA method was benchmarked against existing copy number profile detection methods (i.e., SCOPE (22) and HMMcopy (25)) coupled with classical Hierarchical (27) and K-means (28) clustering methods. Spike-in simulations with pre-defined “true” subclones demonstrated the superior clustering performance of our method. Comparable performance for FLCNA in detecting copy number profile was also observed in simulation studies. Application of FLCNA to a breast cancer dataset successfully clustered subclones based on shared breakpoints of cells in three breast cancer patients. In conclusion, FLCNA was powerful in subclone clustering and CNAs detection for SCS data, especially for data with a large proportion of shared CNAs in a cluster.

## MATERIALS AND METHODS

FLCNA was developed based on a fused lasso model (29) to cluster subclones and simultaneously detect CNAs with scDNA-seq data. Note that though our framework was motivated by the detection of somatic copy number change in tumor tissues, it can also be naturally extended to copy number variation detection as a type of germline genetic variability. For consistency, our framework and evaluation will use the term CNA in describing and evaluating the methods across the manuscript.

### Quality Control and pre-processing of scDNA-seq data

We used scDNA-seq read count data as input. In order to reduce artifacts for copy number detection, a quality control procedure was implemented to remove markers with extreme GC content (< 20% and > 80%) or those with low mappability (< 0.9) (22). Then a two-step median normalization approach (30) was used to further remove the effect of biases from the GC-content and mappability. Basically, the normalized read counts were defined as: 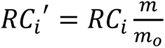, where *RC_i_* was the read counts for marker *i*, *m_o_* was the median read counts of all markers with the same *o* value (where *o*=[the GC-content, mappability]) as the *i*-th marker, and *m* was the overall median read counts of all the markers. We further computed the ratio of the normalized read counts and its sample specific mean, the logarithm transformation of which (i.e., *log2R*) was obtained as the main signal intensities.

### Notations and models

Considering we have *N* cells in total, let ***X*** = (*x_i,j_*)*_P×N_* be the normalized read counts data (i.e., *log2R*), where *x_i,j_* denotes the value of the *i*-th (*i* =1, …, *P*) marker from the *j*-th cell. ***x****_j_* = (*x*_1,*j*_, …, *x_P,j_*)*^T^* is the data vector for the *j*-th sample. The samples sharing common biological characteristics (e.g., genomic variation) belong to a same cluster. Starting from a general *K*-cluster problem with a Gaussian mixture model (GMM), the observations ***x****_j_* are assumed to be independent and are generated from a probability density function 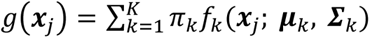, where the “weights” *π_k_′s* (*π_k_* ≥ 0 for each cluster, 1 ≤ *k* ≤ *K* and 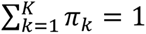) are the mixing proportions. For the *k*-th cluster, *f_k_*(***x****_j_*; ***μ****_k_*, ***Σ_k_***) denotes the Gaussian density function with mean vector ***μ****_k_* = (*μ*_1,*k*_ … *μ_P, k_*)*^T^* and covariance matrix ***Σ****_k_.* ***μ****_k_* denotes the mean genetic intensity of each marker in the *k*-th cluster, and ***Σ****_k_* captures the correlations among the markers. In this study, we assume that the covariance matrix 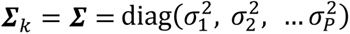 is fixed for all clusters. Parameters including *π_k_*, ***μ****_k_* and ***Σ*** are unknown and need to be estimated.

The problem of copy number profile detection is equivalent to chromosomal breakpoints detection and has been initially explored in the fused lasso model (26). Fused lasso was extended from the classical lasso model and first designed to select variables and penalize the difference of successive features (29). Its ability in identifying and quantifying significant features is closely related to our problem of breakpoints detection on locating significant signals from a wide range of constant signals (26). Utilizing the fused lasso penalty term for change points detection, the penalized log likelihood function in the FLCNA method is given by

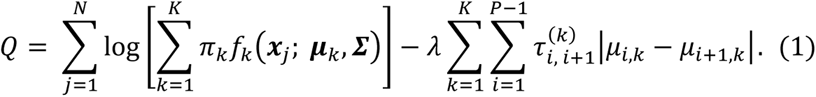

In the second term of Equation (1), a tunning hyperparameter *λ* and pre-defined adaptive weights (31) 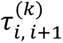 are utilized to shrink the absolute difference of the mean shift values |*μ_i_*_,_*_k_* − *μ_i_*_+1,*k*_| in consecutive markers in the *k*-th cluster, ultimately disclosing change points. The tuning hyperparameter *λ* is used to control the overall number of change points that less change points tend to be generated with larger *λ* value (Figure 1). To shrink each pair of consecutive markers with the same weight, the tunning hyperparameter *λ* is fixed within each cluster. Such strategy may naturally decline the accuracy of penalization, ultimately affecting the clustering performance. To improve the accuracy, an adaptive penalization weight (31) 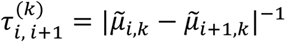 is applied, where 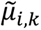 is estimated from the same model without any penalization (*λ* = 0). This adaptive penalization weight is pre-defined to dynamically penalize each pair of successive markers. For example, if there is large difference between 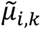 and 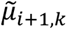 in the model without penalization, a change point is expected to appear between the *i*-th and (*i* + 1)-th marker, and this change point tends to be informative for subclone clustering. In this case, according to 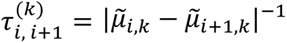, 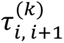 will be small, consequently the difference between *μ_i,k_* and *μ_i_*_+1,_*_k_* in Equation (1) will be lightly penalized, and will be more informative for the subclone clustering. Otherwise, the difference between *μ_i,k_* and *μ_i_*_+1,*k*_ will be heavily penalized and will be less informative for the subclone clustering in the FLCNA method. With the focus on change points detection and subclone identification, our goal is to maximize Equation (1) to estimate the parameter set 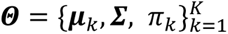.

**Figure 1.**
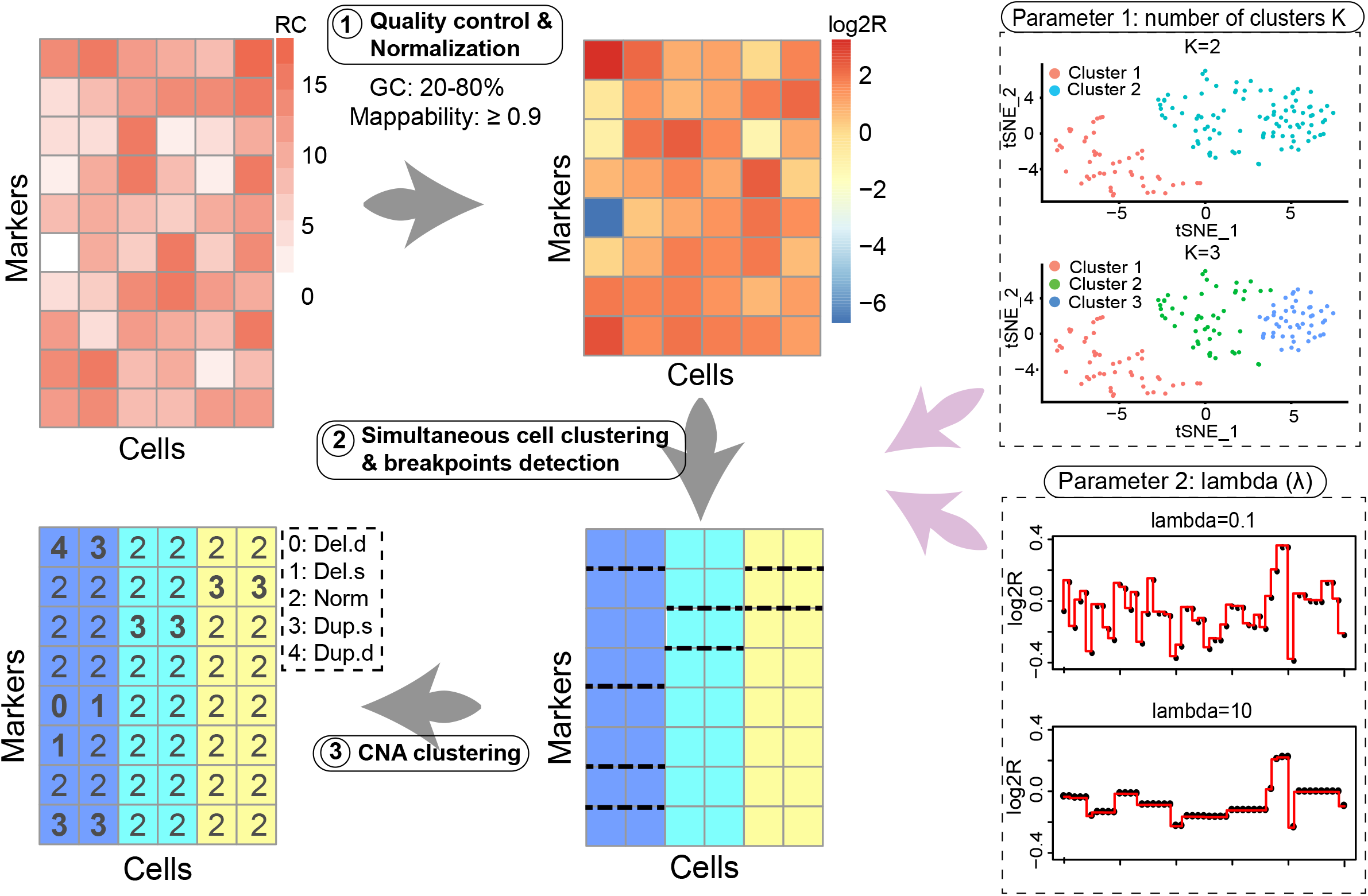
Analysis workflow of FLCNA. FLCNA first implemented a quality control procedure based on GC content and mappability. A two-step median normalization approach was then conducted to sequentially remove the effect of biases. The logarithm transformed ratio of normalized read counts and its sample specific mean (*log2R*) was then used in the main step of the FLCNA method. With *log2R*, we clustered subclones and simultaneously detected shared breakpoints using a Gaussian mixture model (GMM) after adding a fused lasso penalty term. Finally, based on these shared breakpoints in each cluster, segments for each cell were clustered into five different CNA states (Del.d, Del.s, Norm, Dup.s and Dup.d) using *log2R*. There are two hyperparameters in the FLCNA model including the number of cell clusters *K* and a tuning parameter *λ*, which controls the number of breakpoints. Del.d: Deletion of double copies; Del.s: Deletion of a single copy; Norm: Normal/diploid; Dup.s: Duplication of a single copy; Dup.d: Duplication of double copies.

### Parameter estimation using expectation–maximization algorithm

In FLCNA, the parameter set 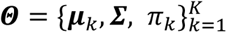 are estimated using expectation–maximization (EM) algorithm (32). We initialize ***Θ*** with parameters estimated from the model without penalty (*λ*=0) and then we update these parameters by alternating between E- and M-steps in the EM algorithm. First, in the M-step, given the starting values of ***Θ***, the probability for the *j*-th sample belonging to the *k*-th cluster is calculated by dividing the density of the *k*-th cluster by the sum of densities from all clusters. Thereafter, an E-step is used to update the estimated values of ***Θ*.** Specifically, the “weight” for the *k*-th cluster *π_k_* and the variance for the *i*-th marker 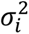 are estimated by taking the first derivative of *Q*(***Θ***) w.r.t. *π_k_* and 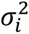, respectively. The estimates of the cluster means 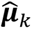 are computed with a local quadratic approximation algorithm (33). These updated ***Θ*** estimation values were iteratively computed between E- and M-steps until convergence. We refer readers to the Supplementary Methods for a thorough description of this part of algorithm. After this step, all the parameters of our interest 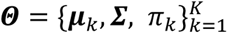 are successfully estimated, based on which copy number profile will be assigned by the estimated cluster means (described below) and cells are clustered according to the estimated cluster weights.

### Copy number profile identification and hyperparameters estimation

With the estimated cluster means (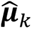), we locate and quantify all the shared change points and identify copy number segments in each cluster. Based on these identified shared segments, we assign the most likely copy number state for each segment in each cell. A GMM-based clustering strategy (34) is implemented for CNA clustering using the normalized read counts data (i.e., *log2R*). Segments sharing similar intensity levels in a cell are identified as the ones with same copy number states. Each segment is classified using a five-state classification scheme with deletion of double copies (Del.d), deletion of a single copy (Del.s), normal/diploid, duplication of a single copy (Dup.s) and duplication of double copies (Dup.d). Besides, there are two hyperparameters to be pre-defined in the FLCNA method, including the number of clusters *K* and the tuning parameter *λ*. To find the optimal values of *K* and *λ*, we use a Bayesian information criterion (BIC) (35) and the clustering model with smallest BIC value is selected as the optimal model.

### Data description

We utilized two publicly available scDNA-seq datasets for illustration and evaluation of FLCNA in simulations and real data analyses. As described below, the BRCA5 dataset was used to mimic real data signals in simulations, and the TNBC dataset was analyzed for real data applications.

The BRCA5 dataset consists of 10,088 cells from a frozen breast tumor tissue that were sequenced using the 10× Genomics platform (https://www.10xgenomics.com), which utilizes microfluidic droplets to barcode cells and performs library construction. We generated a read depth matrix of 28,760 markers and 10,088 cells from BAM files after binning with a 100kb bin size. To save computational time, we randomly selected 220 cells from the dataset and used the read counts from the entire genome to mimic real data in our simulations. More information on the simulation setting is available in the Spike-in simulations section, as described below.

The TNBC dataset consists of data from triple-negative breast cancer patients in the NCBI Sequence Read Archive (SRP114962) (36). TNBC is characterized by extensive intratumor heterogeneity and frequently develops resistance to NAC treatment. Three patients (i.e., KTN126, KTN129, KTN302) were used for our analyses, where tumor cells were only reported in the pre-treatment samples. For each patient, cells were sequenced at two time points (pre- and mid/post-treatment) with 93 cells (46 pre- and 47 post-treatment) in the KTN126 patient, 90 cells (46 pre- and 44 post-treatment) in the KTN129 patient, and 92 cells (47 pre- and 45 mid-treatment) in the KTN302 patient, respectively. For these samples, FASTQ files were generated with Fastq-dump from SRA-Toolkit (37), and then aligned to the NCBI hg19 reference genome and converted to BAM files. Raw read depth of coverage data were generated from the BAM files with a bin size 100kb (22).

### Spike-in simulations

We compared FLCNA to existing copy number profile detection methods, including SCOPE (22) and HMMcopy (25), using spike-in simulations. Since these two methods were only designed for the detection of CNAs without cell clustering, they were followed by two commonly used clustering methods, Hierarchical (27) and K-means (28). The simulation mimicked a scDNA-seq dataset of frozen breast tissue, the BRCA5 dataset (Data description), by randomly selecting 220 cells from the dataset and using SCOPE and HMMcopy to remove genetic regions. Genetic regions with copy number changes detected by either method were excluded from the analysis, and the remaining sequences were treated as copy number-free sequences. Among these cells, 20 cells were randomly selected as reference cells for the SCOPE method, and signals of spiked-in CNAs were added to the remaining cells. To evaluate the robustness of FLCNA, we randomly generated CNAs of varied sizes (super short: 2~5 markers, short: 5~10 markers, medium: 10~20 markers, or long: 20~35 markers) and simulated different numbers of clusters (three or five). We evaluated varied copy number states, including Del.d, Del.s, Dup.s, and Dup.d, respectively. Moreover, data with a mixture of above four different copy number states were generated to mimic real-world data. Because a CNA may not be shared by all the cells in a cluster, different CNA sharing proportions (20%, 40%, 60%, 80%, 100%) were considered. For each cluster, we added 50 CNA segments to the background sequences. Besides, we evaluated the performance on scenarios with different numbers of CNAs among clusters by assigning random numbers of CNAs (20~80) to each cluster. For each of these scenarios, signals were spiked in by multiplying the background depth of coverage by *c*/2, where *c* is a normal random variable following *N*(0.4, 0.1^2^) for Del.d, *N*(1.2, 0.1^2^) for Del.s, *N*(2.8, 0.1^2^) for dup.s and *N*(4.2, 0.1^2^) for dup.d (22). Adjusted Rand Index (ARI) (38) was calculated to evaluate the clustering performance of these methods by comparing the identified clusters of each method to the pre-defined “true” classes. ARI gets close to 1 if clusters identified are completely consistent with the ground truth and close to 0 for random clustering. The performance of CNA detection for these methods was assessed using *F*1 score 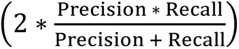 with precision rate defined as 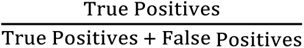 and recall rate defined as 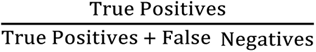.

### FLCNA application to the TNBC dataset for breast cancer subcloning

With the dataset introduced previously, FLCNA was also applied to the TNBC dataset of breast cancer with three unrelated patients (i.e., KTN126, KTN129, KTN302) who have been treated with NAC. We identified shared CNAs using FLCNA and mapped them to 575 significant genes from the genome-wide association studies (GWAS) with breast cancer curated in the NHGRI-EBI GWAS Catalog (39). Pathway and network analyses were conducted for these genes; Gene Set Enrichment Analysis (GSEA) (40) was conducted with enrichment of Kyoto Encyclopedia of Genes and Genomes (KEGG) (41). Further, the summary statistics from GSEA were used to generate connection networks for these three patients using the Enrichment Map implemented in Cytoscape (42).

## RESULTS

To capture the biological heterogeneity between potential subclones, we developed the FLCNA method based on a fused lasso model, which can simultaneously identify subclones and detect breakpoints in scDNA-seq data. The framework of the FLCNA method is summarized and illustrated in Figure 1. First, quality control and normalization procedures were used for pre-processing the datasets. Subclone clustering and breakpoints detection were achieved simultaneously using GMM combined with a fused lasso penalty term. Finally, based on these shared breakpoints in each cluster, candidate CNA segments for each cell were clustered into different CNA states using a GMM-based clustering strategy. Through extensive simulations, we evaluated the performance of FLCNA in clustering subclones and detecting CNAs. A real dataset of breast cancer (i.e., TNBC) was also utilized to demonstrate the application of FLCNA in scDNA-seq data.

### Evaluation of FLCNA via spike-in simulations

We first conducted spike-in simulations to assess the clustering performance of FLCNA, compared to two other copy number estimation methods (SCOPE and HMMcopy) coupled with different clustering methods (Hierarchical or K-means). Using a mixture of four different copy number states (Del.d, Del.s, Dup.s and Dup.d), we found that FLCNA outperformed the other methods in all scenarios with different numbers of clusters and CNAs in a cluster (Figure 2 and Supplementary Figure S1–S2). Specifically, with five pre-defined clusters and a fixed number of CNAs (i.e., 50) in each cluster, FLCNA’s clustering performance was incrementally improved as the CNA sharing proportion increased, and it generally presented better ARI than the other methods, especially with larger CNA sharing proportion (> 40%) (Figure 2 and Supplementary Table S1). However, when the CNA sharing proportion decreased to 20%, almost all methods failed to provide desirable clustering performance (ARI < 0.50).

**Figure 2.**
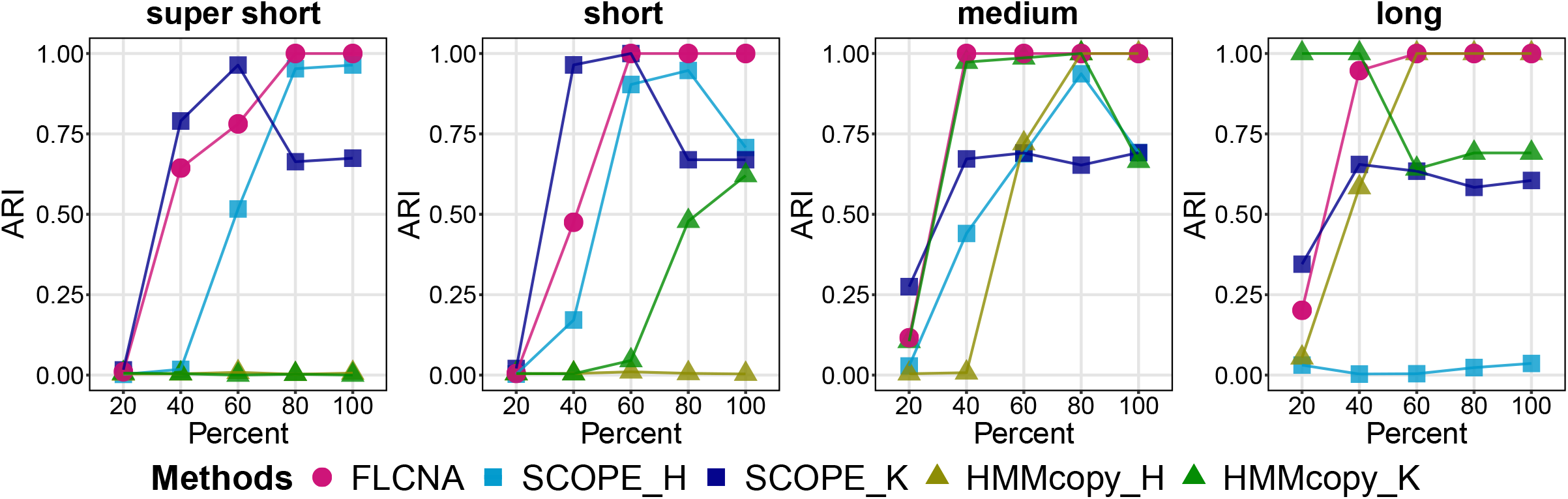
Assessment of FLCNA using simulation data with five clusters and mixed CNA states. Clustering results from FLCNA were compared to existing methods (i.e., SCOPE and HMMcopy) coupled with different clustering methods. For each of five clusters, we added signals of 50 CNA segments to the background signals with varied lengths (super short: 2~5 markers, short: 5~10 markers, medium: 10~20 markers, and long: 20~35 markers) and varied CNA proportions (20%, 40%, 60%, 80%, 100%), respectively. Signals of mixed CNA states (i.e., Del.d, Del.s, Norm, Dup.s and Dup.d) were spiked in. ARI: Adjusted Rand Index; SCOPE_H: SCOPE_Hierarchical; SCOPE_K: SCOPE_K-means; HMMcopy_H: HMMcopy_Hierarchical; HMMcopy_K: HMMcopy_K-means.

Moreover, the overall clustering performance of all methods was improved in scenarios with three pre-defined clusters (Supplementary Figure S1 and Table S2). For example, for super short CNAs shared by 60% of samples, ARI of FLCNA improved from 0.781 in the scenario of five pre-clusters (Supplementary Table S1) to 0.985 in that of three pre-clusters (Supplementary Table S2). This improvement was likely due to the increased sample size in each cluster given a fixed total sample size. Comparing to the scenario with a fixed number of shared CNAs in each cluster, it was more challenging for all methods to identify subclones when varied numbers of shared CNAs presented, due to fewer shared CNAs generated in some clusters by randomness (Supplementary Figure S2 and Table S3). For samples with a single type of copy number state (i.e., Del.d, Del.s, Dup.s, Dup.d), FLCNA still outperformed other methods in detecting deletions or duplications (Supplementary Figure S3–S5 and Table S1-S3). All methods provided higher ARI in deletions than duplications, and in double copy changes than single copy, due to the stronger signals presented in the intensities for the former than the latter cases. Overall, FLCNA was the best method in detecting commonly shared CNAs.

We also evaluated the accuracy of FLCNA in CNA detection by comparing it to SCOPE and HMMcopy (Figure 3 and Supplementary Figure S6–S10, Table S4-S6). FLCNA demonstrated improved accuracy in identifying CNAs as the proportion of shared CNAs increased. For example, with a fixed number of CNAs (i.e., 50) shared by all samples in each of five clusters, FLCNA had an *F*1 score of 0.896, while SCOPE only had a score of 0.435 and HMMcopy had a score of 0.783 (Figure 3 and Supplementary Table 4). In general, FLCNA outperformed the other methods in detecting super short and short CNAs.

**Figure 3.**
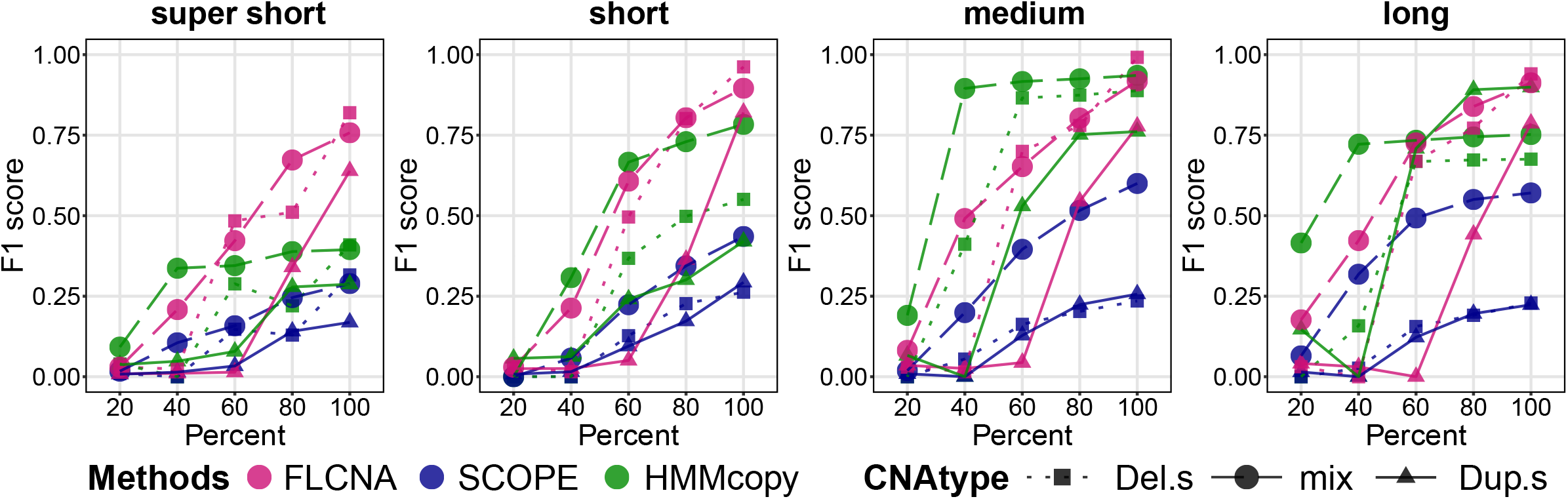
Assessment of FLCNA to detect CNAs using simulation data with five clusters. CNA calls were generated by FLCNA, SCOPE and HMMcopy, respectively. For each of five clusters, we added signals of 50 CNA segments to the background signals with varied lengths (super short: 2~5 markers, short: 5~10 markers, medium: 10~20 markers, and long: 20~35 markers) and varied CNA proportions (20%, 40%, 60%, 80%, 100%), respectively. Deletion of a single copy (Del.s), mixed CNA states (mix) and duplication of a single copy (Dup.s) were spiked in separately. *F*1 score was utilized to evaluate the performance of CNA detection for each method.

In summary, FLCNA showed overall great performance in clustering subclones with copy number changes shared by a large proportion of samples within a cluster. In copy number profile detection, FLCNA presented its advantage in detecting short CNAs shared by a relatively larger proportion of samples within a cluster.

### Application to the TNBC breast cancer single cell study

FLCNA was also applied to a single cell study of three breast cancer patients, resulting in the identification of three clusters in the KTN126 patient and two in the other two patients (Figure 4 and Supplementary Figure S11–S12). In particular, for the KTN126 patient, 64 cells (17 cells with pre-treatment and 47 cells with post-treatment) were clustered in cluster A, with 9 cells in cluster B and 20 cells in cluster C, all from the pre-treatment group. Similar copy number profile patterns were observed within clusters. Interestingly, clusters B and C, which included more pre-treatment samples, showed higher variation in genetic intensities, indicating that some copy number aberrations were treatment-specific. A more interesting and feasible interpretation is that the treatment of NAC may lead to extinction of tumor cells with these copy number changes in this patient. Similar patterns were observed in the other two patients (Supplementary Figure S11–S12), and the locations of these treatment-specific copy number changes were consistent across patients KTN129 and KTN302.

**Figure 4.**
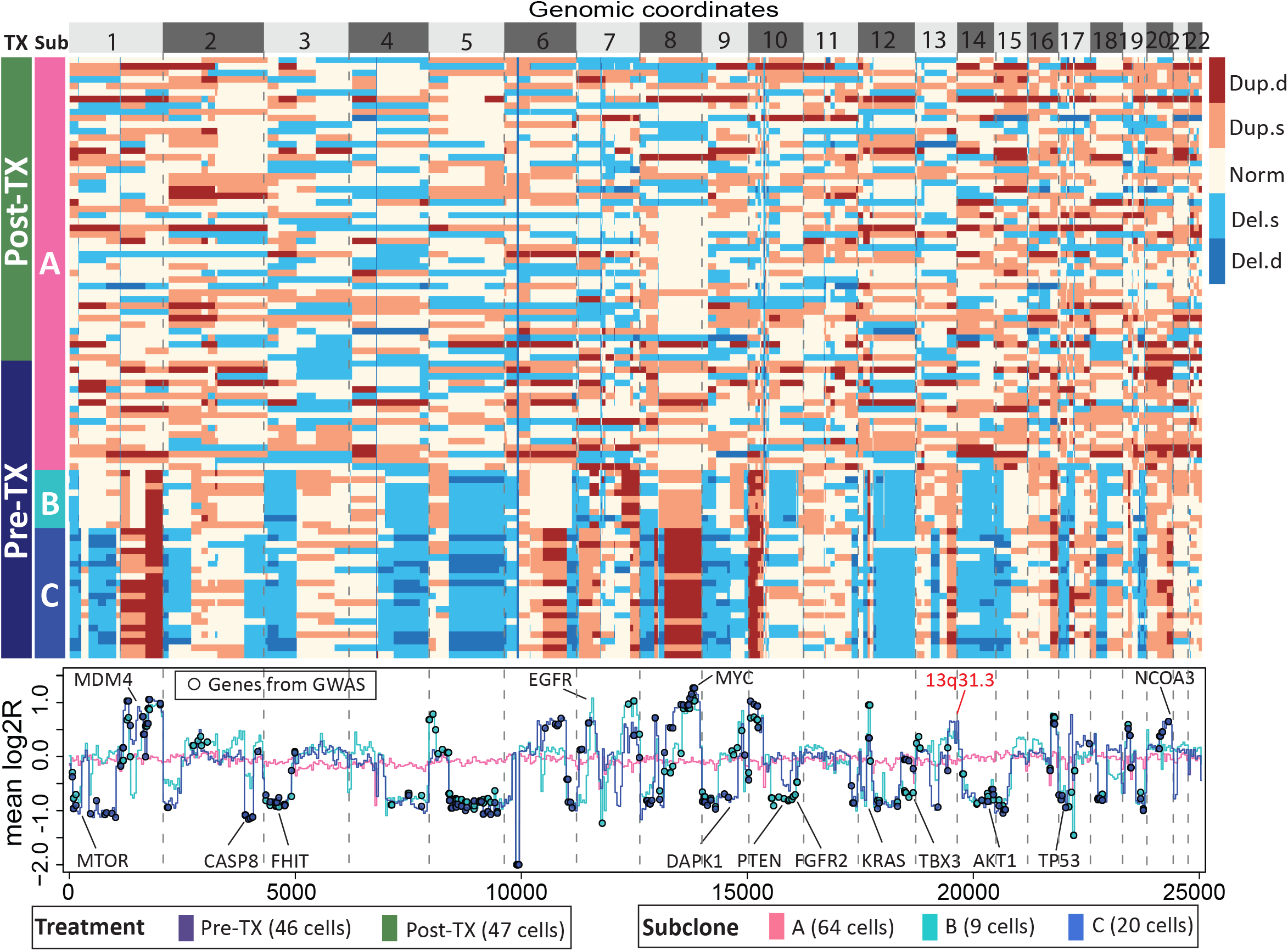
Subclone clustering of the KTN126 patient using FLCNA. Cell clusters and copy number profile with different CNA states (Del.d, Del.s, Norm, Dup.s and Dup.d) were generated using FLCNA. Mean log2R were provided for each cluster. Shared CNAs identified using FLCNA were matched to significant genes from the genome-wide association studies (GWAS) in the NHGRI-EBI GWAS Catalog. Del.d: Deletion of double copies; Del.s: Deletion of a single copy; Norm: Normal/diploid; Dup.s: Duplication of a single copy; Dup.d: Duplication of double copies; log2R: Logarithm transformation of ratio between normalized read counts and its sample specific mean; Pre-TX: pre-treatment; Post-TX: post-treatment.

FLCNA identified 264, 154 and 156 shared CNAs in KTN126, KTN129 and KTN302 patients, respectively. Consensus CNA located genes (e.g., *DAPK1*, *TBX3*, *NCOA3*, *KRAS*) among patients implied the shared common evolutionary path of the tumor cells. Mutations in these genes have been shown to play importance roles in the evolution (*DAPK1* (43), *TBX3* (44)) or the therapy (*NCOA3* (45), *KRAS* (46)) of breast cancer. The shared CNAs were mapped to 436 out of 575 breast cancer risk-associated genes identified from existing GWAS. These mapped genes were enriched in pathways related to cancer, hormones, immunity, and epithelial-mesenchymal transition (EMT) (Supplementary Figure S13). Most pathways were shared by all three patients and consistent with findings from existing studies on breast cancer. For instance, PATHWAYS_IN_CANCER was found to be hyperactivated in the human tumor tissue (47). EMT (e.g., ADHERENS_JUNCTION(48)) and immune (e.g., TOLL_LIKE _RECEPTOR(49)) related pathways were associated with the invasion and metastasis of tumor cells. Various hormones also played importance roles in the occurrence and progression of breast cancer (50). We also observed a novel CNA in 13q31.3 (e.g., *LINC01040*) among all three patients, with duplications in KTN126 and KTN129 patients and deletions in the KTN302 patient. This discovery of a new CNA may provide new insights in understanding the progression of breast cancer.

To evaluate the heterogeneity of CNAs among cells, the sharing proportion of CNAs was evaluated by cluster. The mean sharing proportion was 30.3% in the KTN126 patient (18.6% in cluster A, 36.4% in cluster B, and 34.7% in cluster C), 25.7% in the KTN129 patient, and 26.4% in the KTN302 patient (Supplementary Figure S14). Overall, our method detected CNAs with a wide range of length, varying from 2 to 1,000 bins (Supplementary Figure S15). For computational speed, with 93 cells from the KTN126 patient (Supplementary Table S7), FLCNA was much faster (1.2 hours) than SCOPE (10.5 hours) using a high-performance cluster with 12GB RAM, whereas HMMcopy took 0.5 hours.

## DISCUSSION

The importance of copy number change in modulating human disease is increasingly being recognized. scDNA-seq enables researchers to profile diseases and biological processes at the single-cell level. Accurate detection of CNAs with scDNA-seq data is crucial for identifying copy number profiles at a single-cell resolution, ultimately leading to a better understanding of how tumor lineages evolve (51, 52). Numerous scDNA-seq studies (18, 53, 54) have used CNAs to characterize tumor subclones, and have found that most tumors contain multiple subclonal lineages. In our study, we developed FLCNA based on the fused lasso model for copy number estimation and simultaneous subclone clustering. Our simulations demonstrated the desirable performance of our method in clustering subclones and estimating copy number with scDNA-seq data, especially for data with a large sharing proportion of CNAs within a cluster.

Recent advances in NGS and SCS technologies have provided emerging tools for computational methods to infer subclones with different genomic variants, including single nucleotide alteration (SNAs) and CNAs. SAPPH (55) estimated CNAs and infer tumor subclone proportion from paired tumor-normal data. The SCClone (56) and SiCloneFit (57) methods clustered subclones using single cell SNAs data to reconstruct the clonal populations and the evolutionary relationship between the clones. Elyanow *et al*. (58) overcame the challenges in inferring CNAs from RNA-sequencing data by utilizing spatial information of a small group of cells to help identify CNAs and the spatial distribution of clones within a tumor sample. A major distinguishing feature of our method is its simultaneous clustering of cells and detection of CNAs, which solves the problem of declined clustering performance resulting from falsely identified variants in the stage of variants estimation.

The superior clustering performance of FLCNA has been demonstrated in extensive simulations. Our method was developed specifically for scDNA-seq with a large proportion of shared CNAs, which are more prevalent in single cell data than in bulk sequencing data or combined-cell samples (59). Consistently, our spike-in simulations provided powerful clustering results from FLCNA for the samples with a large proportion of shared CNAs. Using a whole genome dataset with breast cancer, FLCNA successfully clustered subclones and provided clearly different genomic variation patterns in different clusters. With real dataset, we also found that treatment of NAC may lead to extinction of tumor cells despite of the possible bias brought by the fact that partial of the post-treatment samples were collected adjacent to normal tissues. In addition to prediction accuracy, the high dimensionality nature of scDNA-seq datasets desires for high computational efficiency. Even with 100kb in the binning size instead of 500kb as used in SCOPE, our method remained a high computational speed, meanwhile inclusively allowing the accurate detection of short CNAs.

As with any study, there are limitations in our study. First, only shared breakpoints identified in a cluster were used to estimate the underlying copy number profile for each cell, which might not be desirable when the goal is to identify CNAs at the single cell level with a relatively low sharing proportion in this cluster. Additionally, errors due to doublets in single cell sequencing data, where ≥ 2 cells are accidently mixed for sequencing (60), were not considered in our modeling. Doublets have been found to affect the accuracy of copy number profile estimation from previous studies (60). However, since only shared breakpoints were utilized for CNA detection in our study, which may neutralize the bias arising from the existence of doublets since the effect of a few doublets might be diluted by other majority cells from the same cluster. As a result, breakpoints can still be accurately detected.

In conclusion, our FLCNA method sheds new light on the methodology development of CNA detection and subclone identification in single-cell sequencing data using a novel statistical learning strategy. Modern sequencing technologies are continually emerging with multi-omics, spatial or temporal capabilities, which can facilitate better understanding of how subclonal lineages associate with cellular phenotypes, invasiveness and treatment responses (61). Our method has the potential to extend to other data types, such as scRNA-seq, and integrate different dimensions of sequencing information (e.g., gene expression, spatial information) to improve the identification of subclones and benefit research in cancer outcome-related targeted therapy. For example, we can incorporate the spatial information into our model for more comprehensive inference of subclones with scDNA-seq data, as cells located nearby are more likely to share similar genetic patterns and consequently tend to reside within same subclones (58).

## DATA AVAILABILITY

The BRCA5 dataset is available in the 10× Genomics platform (https://www.10xgenomics.com). The TNBC dataset is available in NCBI Sequence Read Archive with accession code SRP114962. FLCNA source code and documentation are publicly available at https://github.com/FeifeiXiaoUSC/FLCNA.

## ACKNOWLEDGEMENTS

We thank the reviewers in advance for their thoughtful and insightful comments.

## Supplementary Methods

### Parameter estimation using expectation–maximization algorithm

In FLCNA, the parameter set 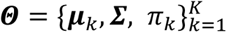 are estimated using an expectation–maximization (EM) algorithm. let Δ*_j,k_* be an indicator function of the hidden cluster information for ***x****_j_*, Δ*_j,k_*= 1 if ***x****_j_* is from the *k*-th cluster, and Δ*_j,k_*= 0 otherwise. Assuming Δ*_j,k_* is unobserved, the penalized log-likelihood function for the complete data will be given by

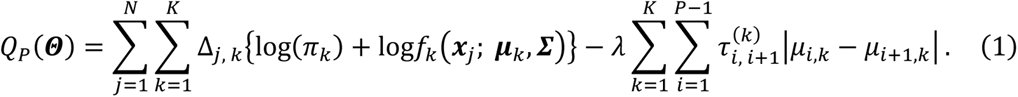

With Eq. (1), the parameter set 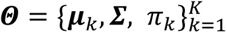 can be estimated with the EM algorithm by the following iterative procedure. The EM algorithm iterates between E-step and M-step, and produces a sequence of estimates 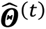, *t* = 0, 1, 2, ….

1) Initialization: We first estimate the starting values 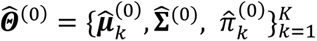 using model without penalty (*λ*=0).

2) Iteration:

E-step:

We start with the E-step given the current parameter estimates 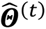. In this step, we calculate the probability for sample *j* belongs to *k*-th cluster with

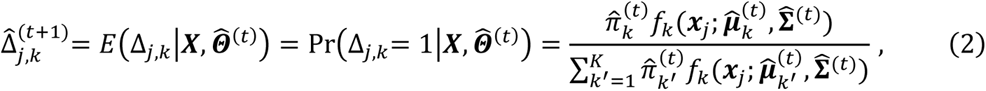

where the numerator is the density for *j*-th sample to be clustered into *k*-th cluster, and the denominator is the sum of densities for *j*-th sample to be clustered into *K* different clusters. Then Eq. (2) will be plugged it into the Eq. (1) about *Q_P_*(***Θ***) to estimate other parameters, including the cluster “weight” *π_k_*, the variance for *i*-th marker 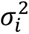 and cluster mean ***μ***.

M-Step:

Given 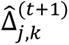 and 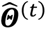, the goal of M-step is to update parameter set 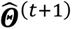 by maximizing the loglikelihood function 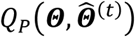. Specifically, the estimate of “weights” *π_k_′s* can be easily updated by taking the first derivative of 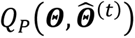 w.r.t. *π_k_* with

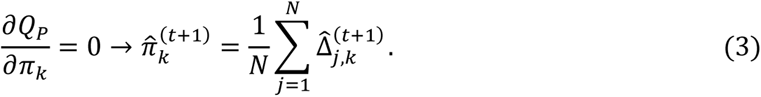

Given 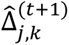, 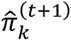 and 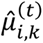, we can update the estimate of variance for *i*-th marker 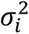 by taking the first derivative of 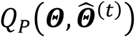 w.r.t. 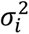 with

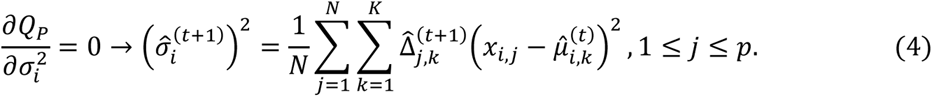

Given 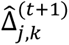, 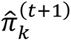 and 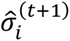, according to Eq. (1), after some transformation, we can update the estimates of mean values 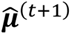 with

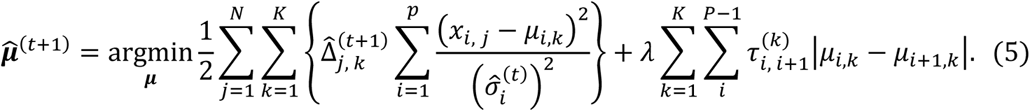

Eq. (5) cannot be solved directly with close form, but 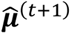 can be estimated using a local quadratic approximation (LQA) algorithm, which will be discussed in detail next.

### Estimation of 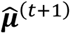 using local quadratic approximation

According to LQA, we can approximate

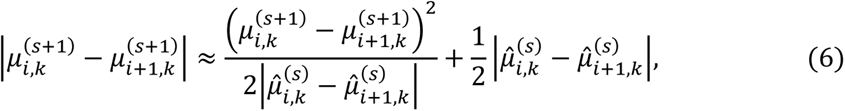

where *s* is the iteration index used to denote iterations of the LQA within the M-step (different from iteration index *t* in the EM algorithm), and 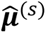 are the estimates from the previous iteration. Thus, the minimization problem in Eq. (5) has been converted into a generalized quadratic problem which has close form solution. Notably, Eq. (5) can be decomposed into *K* separate minimization problems. For example, for each *k*, we can solve (iteratively over *s*)

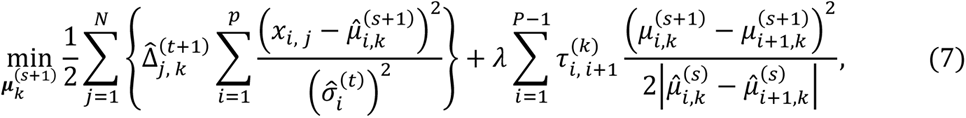

with close form. To solve Eq. (7), we need to transfer it into matrix form first. Let

- 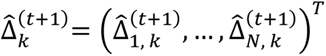 be the estimated latent variable for the *k*-cluster from E-step in the EM algorithm.
- ***J****_N×_*_1_ = (1, …, 1)*^T^* is a matrix with all elements to be 1.
- 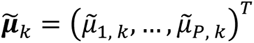 is the pre-defined mean vector for the *k*-th cluster where 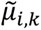 is estimated from the model without any penalization (*λ* = 0).
- 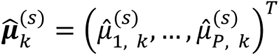 is the estimate of mean vector for the *k*-th cluster from previous iteration in the EM algorithm.
- 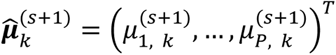 is the estimate of our interest which is the mean vector for the *k*-th cluster.
- 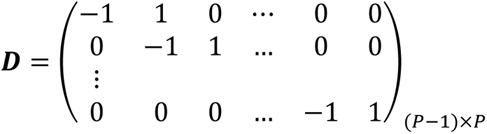 is a matrix introduced to calculate the difference of mean values for each pair of consecutive markers in a cluster.

Then Eq. (1) can also be given with

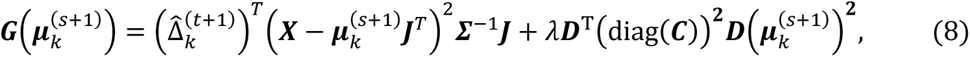

where 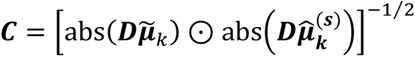.

Thus, we can easily find the solution for the quadratic equation of 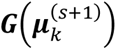 with respect to 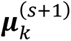,

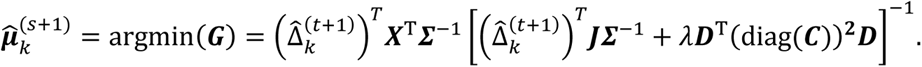

### Model selection

There are two hyperparameters to be pre-defined in the FLCNA method, including the number of clusters *K* and the tuning parameter *λ*. To find the optimal values of *K* and *λ*, we use a Bayesian information criterion (BIC), defined by

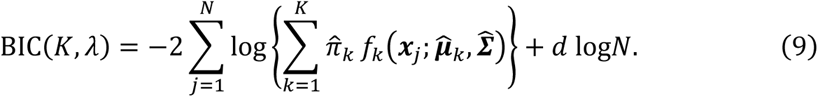

The degrees of freedom 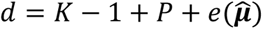, where 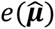 is the number of distinct nonzero elements in 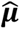 and was used to adjust the number of breakpoints in degree of freedom. For each pair of parameter values (*K*, *λ*), the clustering model with smallest BIC value is selected as the optimal model and the corresponding parameters are estimated.

## Supplementary Figures

**Supplementary Figure 1.**
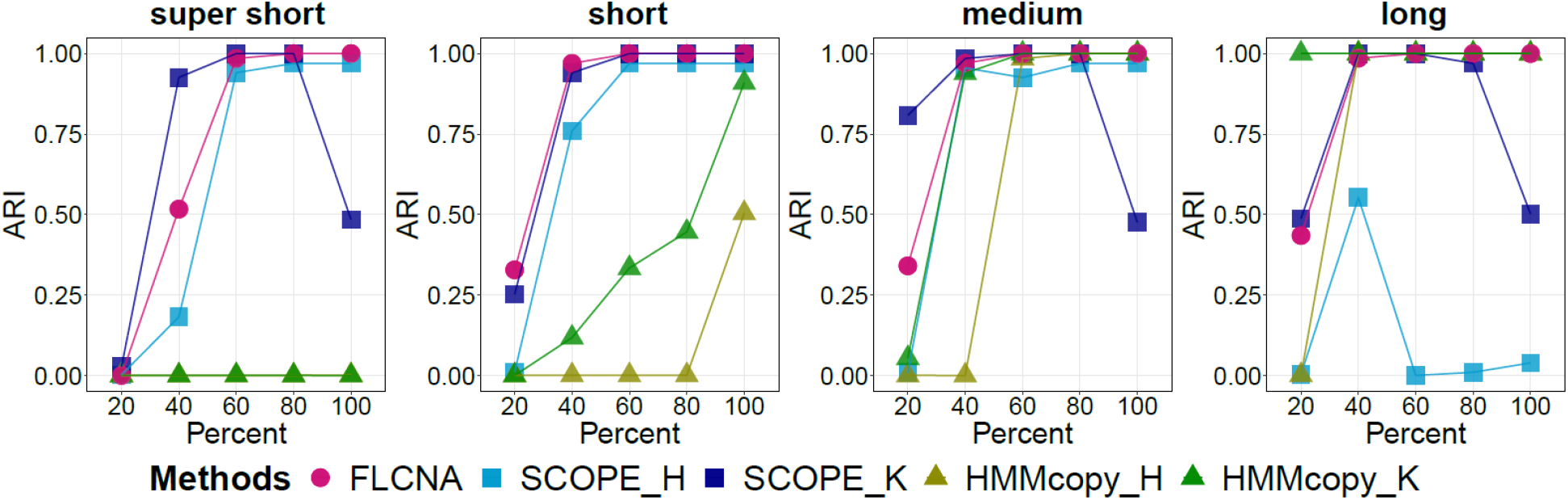
Assessment of FLCNA using simulation data with three clusters and mixed CNA states. Clustering results from FLCNA were compared to existing methods (i.e., SCOPE and HMMcopy) coupled with different clustering methods. For each of three clusters, we added signals of 50 CNA segments to the background signals with varied lengths (super short: 2~5 markers, short: 5~10 markers, medium: 10~20 markers, and long: 20~35 markers) and varied CNA proportions (20%, 40%, 60%, 80%, 100%), respectively. Signals of mixed CNA states (i.e., Del.d, Del.s, Norm, Dup.s and Dup.d) were spiked in. ARI: Adjusted Rand Index; SCOPE_H: SCOPE_Hierarchical; SCOPE_K: SCOPE_K-means; HMMcopy_H: HMMcopy_Hierarchical; HMMcopy_K: HMMcopy_K-means.

**Supplementary Figure 2.**
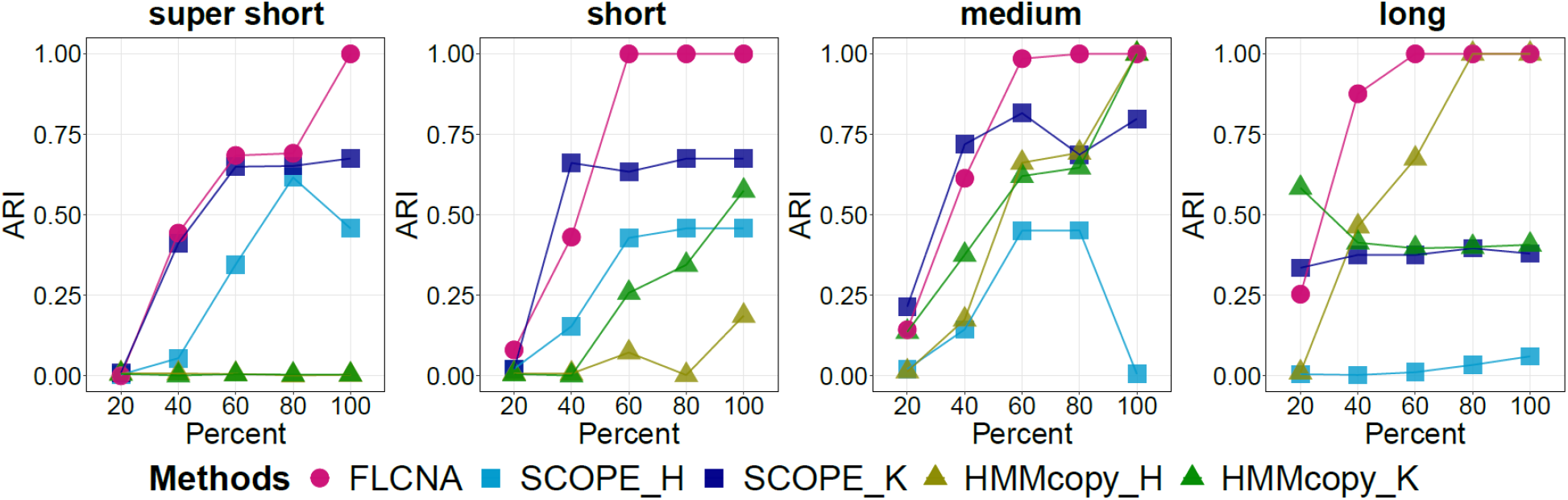
Assessment of FLCNA using simulation data with five clusters, varied numbers of CNAs and mixed CNA states. Clustering results from FLCNA were compared to existing methods (i.e., SCOPE and HMMcopy) coupled with different clustering methods. For each of five clusters, we added signals of varied numbers of CNA segments (20~80) to the background signals with varied lengths (super short: 2~5 markers, short: 5~10 markers, medium: 10~20 markers, and long: 20~35 markers) and varied CNA proportions (20%, 40%, 60%, 80%, 100%), respectively. Signals of mixed CNA states (i.e., Del.d, Del.s, Norm, Dup.s and Dup.d) were spiked in. ARI: Adjusted Rand Index; SCOPE_H: SCOPE_Hierarchical; SCOPE_K: SCOPE_K-means; HMMcopy_H: HMMcopy_Hierarchical; HMMcopy_K: HMMcopy_K-means.

**Supplementary Figure 3.**
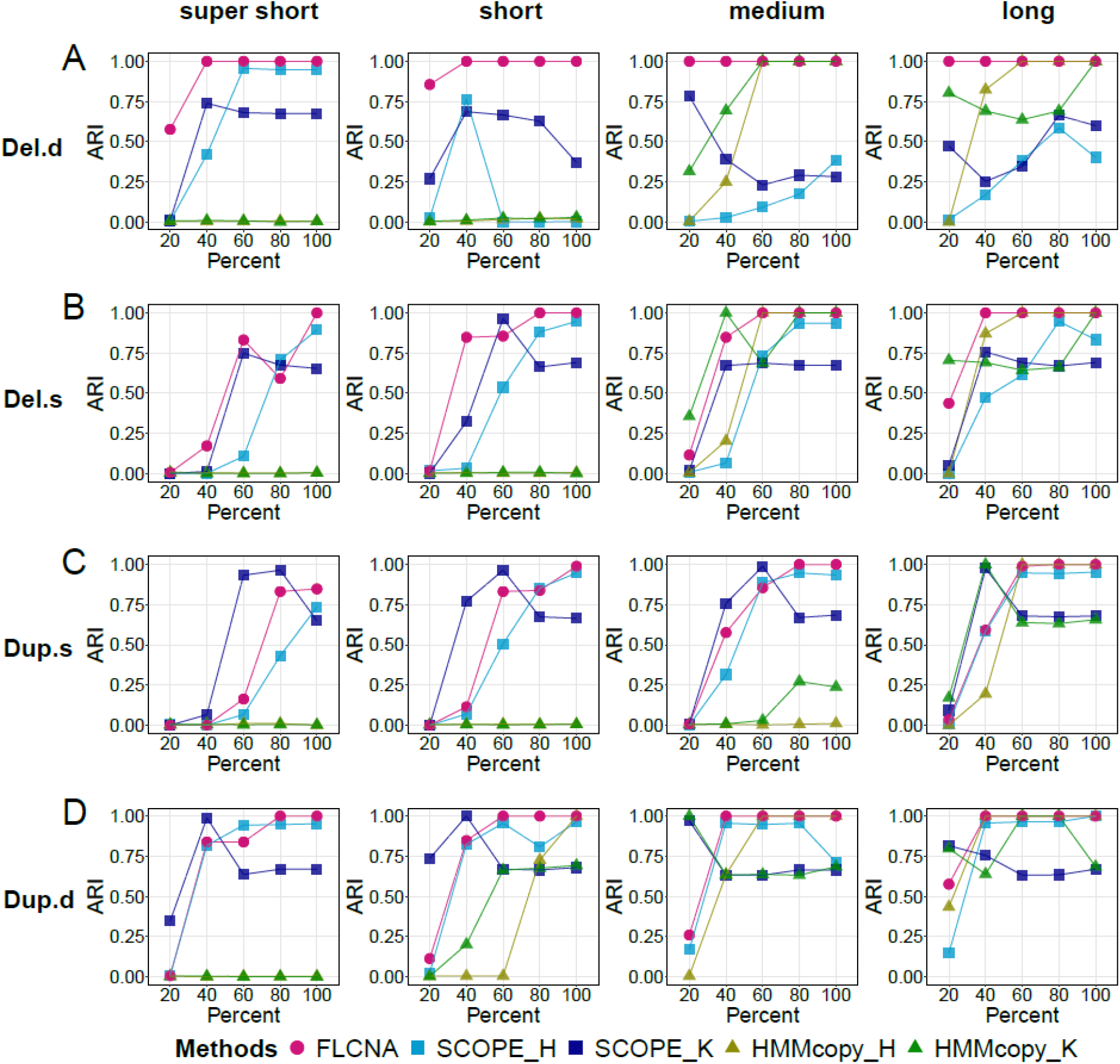
Assessment of FLCNA using simulation data with five clusters and a single type of CNA state. Clustering results from FLCNA were compared to existing methods (i.e., SCOPE and HMMcopy) coupled with different clustering methods. For each of five clusters, we added signals of 50 CNA segments to the background signals with varied lengths (super short: 2~5 markers, short: 5~10 markers, medium: 10~20 markers, and long: 20~35 markers) and varied CNA proportions (20%, 40%, 60%, 80%, 100%), respectively. Signals of Del.d (A), Del.s (B), Dup.s (C) and Dup.d (D) were spiked in separately. ARI: Adjusted Rand Index; Del.d: Deletion of double copies; Del.s: Deletion of a single copy; Dup.s: Duplication of a single copy; Dup.d: Duplication of double copies; SCOPE_H: SCOPE_Hierarchical; SCOPE_K: SCOPE_K-means; HMMcopy_H: HMMcopy_Hierarchical; HMMcopy_K: HMMcopy_K-means.

**Supplementary Figure 4.**
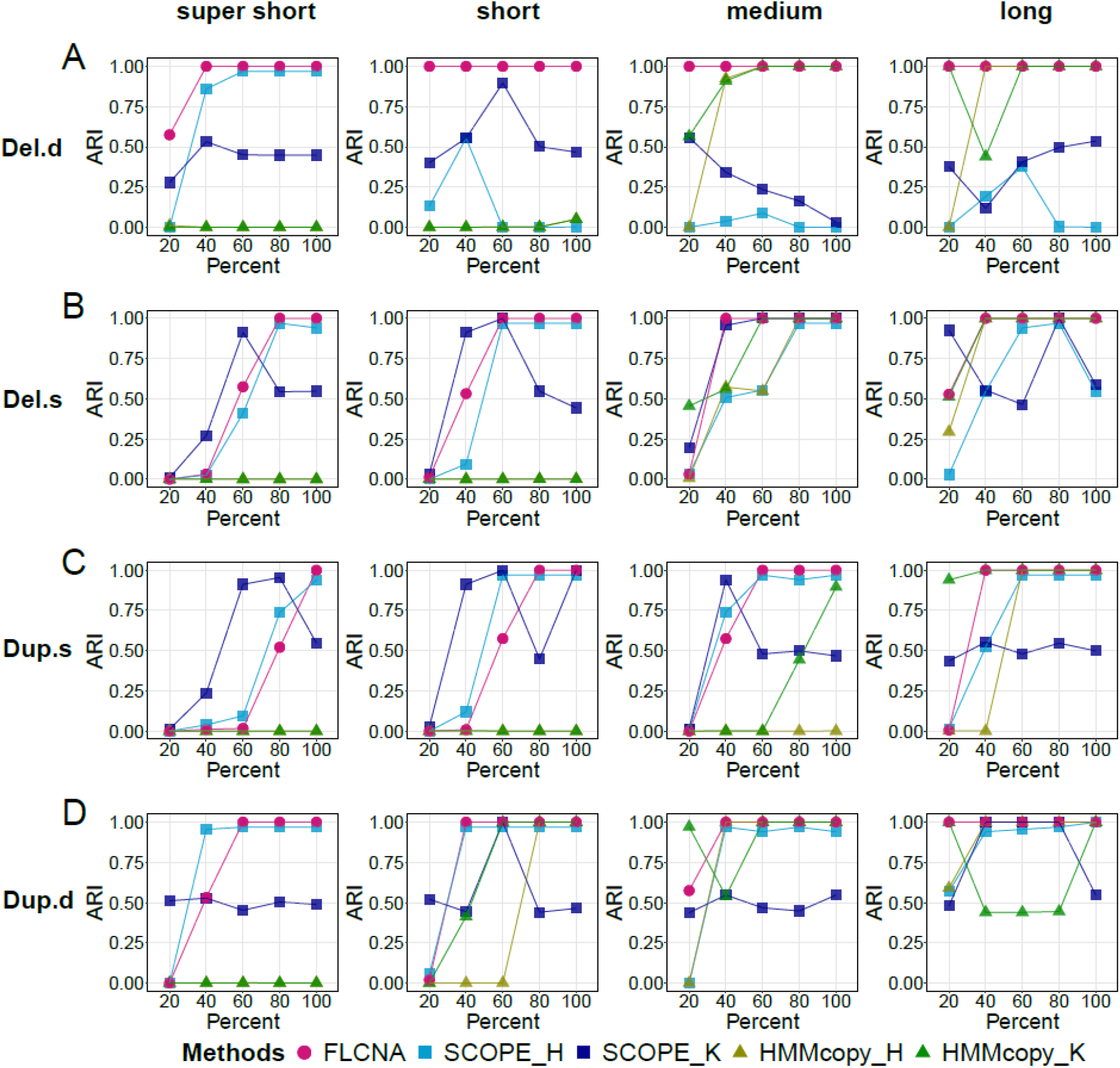
Assessment of FLCNA using simulation data with three clusters and a single type of CNA state. Clustering results from FLCNA were compared to existing methods (i.e., SCOPE and HMMcopy) coupled with different clustering methods. For each of three clusters, we added signals of 50 CNA segments to the background signals with varied lengths (super short: 2~5 markers, short: 5~10 markers, medium: 10~20 markers, and long: 20~35 markers) and varied CNA proportions (20%, 40%, 60%, 80%, 100%), respectively. Signals of Del.d (A), Del.s (B), Dup.s (C) and Dup.d (D) were spiked in separately. ARI: Adjusted Rand Index; Del.d: Deletion of double copies; Del.s: Deletion of a single copy; Dup.s: Duplication of a single copy; Dup.d: Duplication of double copies; SCOPE_H: SCOPE_Hierarchical; SCOPE_K: SCOPE_K-means; HMMcopy_H: HMMcopy_Hierarchical; HMMcopy_K: HMMcopy_K-means.

**Supplementary Figure 5.**
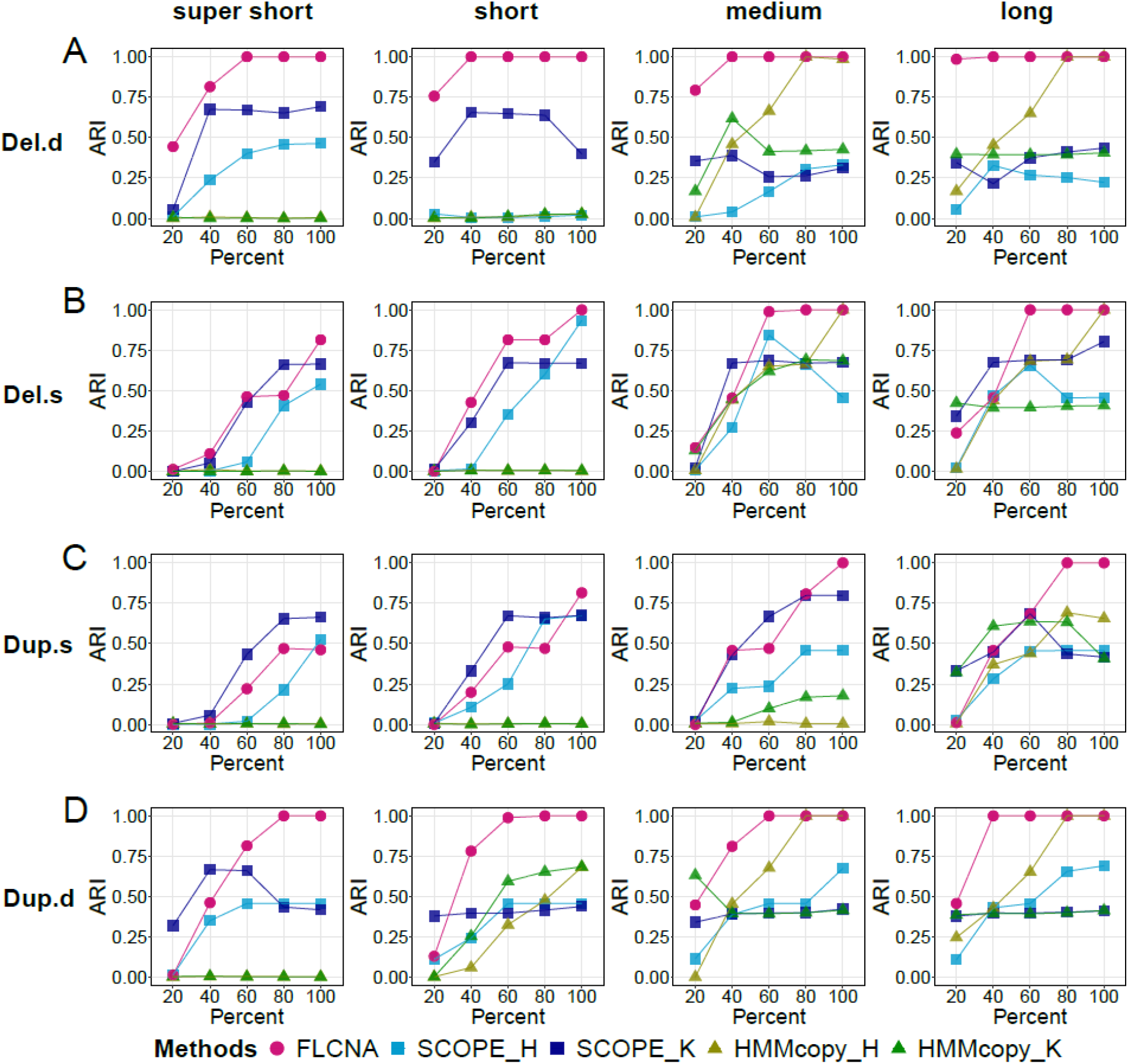
Assessment of FLCNA using simulation data with five clusters, varied numbers of CNAs and a single type of CNA state. Clustering results from FLCNA were compared to existing methods (i.e., SCOPE and HMMcopy) coupled with different clustering methods. For each of five clusters, we added signals of varied numbers of CNA segments (20~80) to the background signals with varied lengths (super short: 2~5 markers, short: 5~10 markers, medium: 10~20 markers, and long: 20~35 markers) and varied CNA proportions (20%, 40%, 60%, 80%, 100%), respectively. Signals of Del.d (A), Del.s (B), Dup.s (C) and Dup.d (D) were spiked in separately. ARI: Adjusted Rand Index; Del.d: Deletion of double copies; Del.s: Deletion of a single copy; Dup.s: Duplication of a single copy; Dup.d: Duplication of double copies; SCOPE_H: SCOPE_Hierarchical; SCOPE_K: SCOPE_K-means; HMMcopy_H: HMMcopy_Hierarchical; HMMcopy_K: HMMcopy_K-means.

**Supplementary Figure 6.**
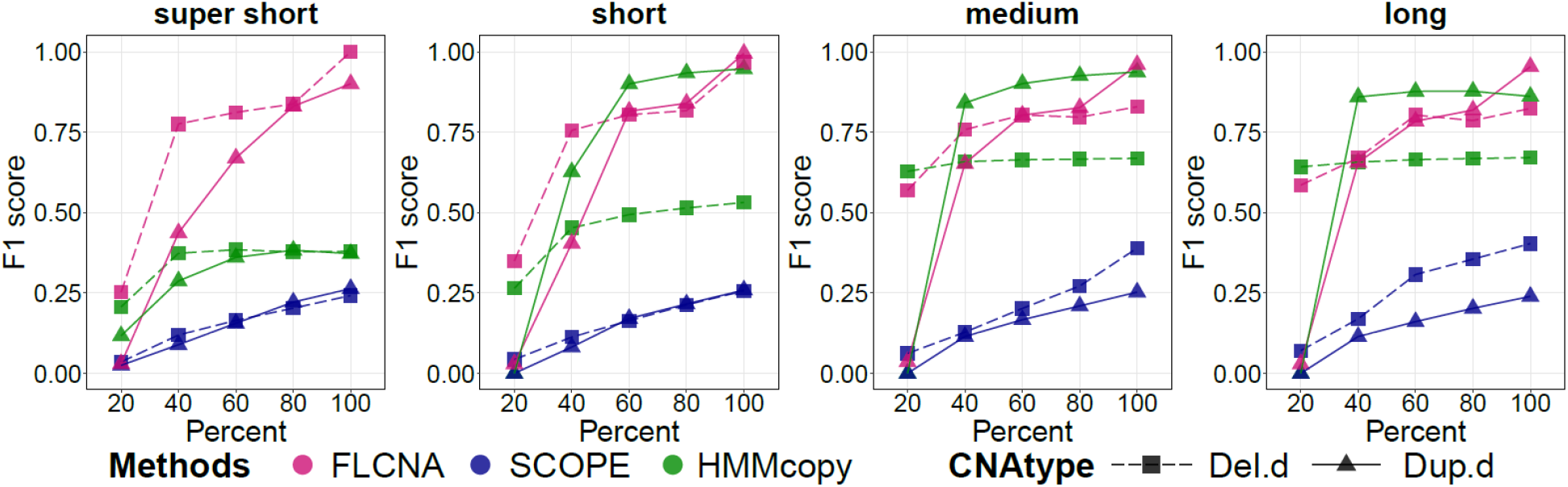
Assessment of FLCNA to detect CNAs using simulation data with five clusters and aberration of double copies. CNA calls were generated by FLCNA, SCOPE and HMMcopy, respectively. For each of five clusters, we added signals of 50 CNA segments to the background signals with varied lengths (super short: 2~5 markers, short: 5~10 markers, medium: 10~20 markers, and long: 20~35 markers) and varied CNA proportions (20%, 40%, 60%, 80%, 100%), respectively. Deletion of double copies (Del.d) and duplication of double copies (Dup.d) were spiked in separately. *F*1 score was utilized to evaluate the performance of CNA detection for each method.

**Supplementary Figure 7.**
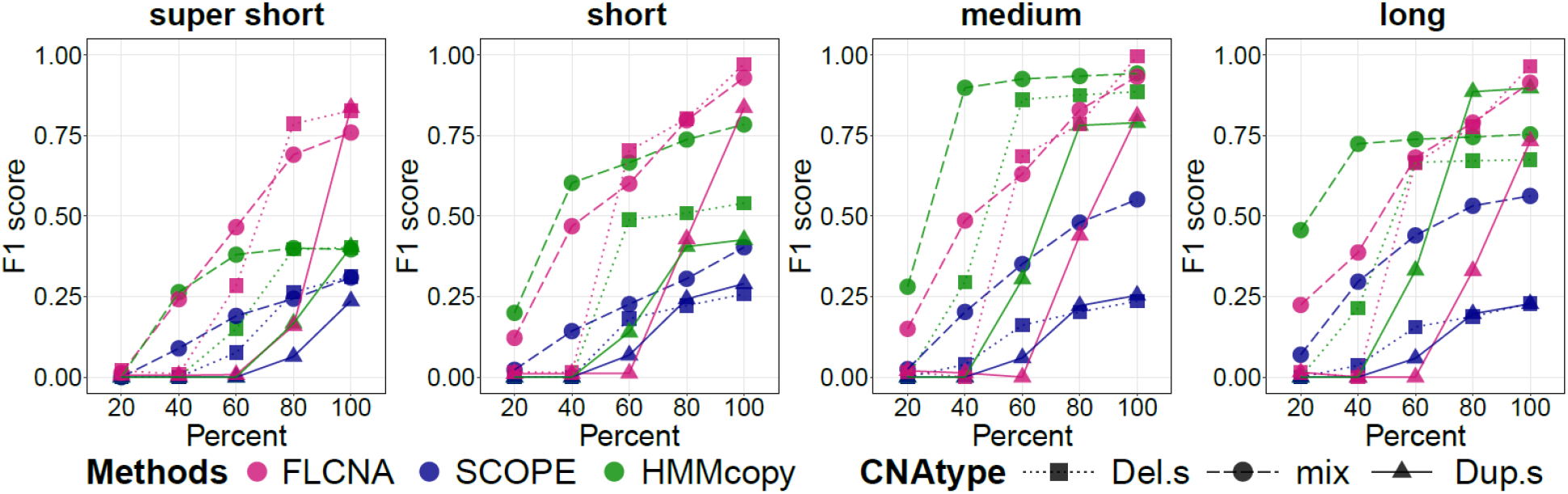
Assessment of FLCNA to detect CNAs using simulation data with three clusters. CNA calls were generated by FLCNA, SCOPE and HMMcopy, respectively. For each of three clusters, we added signals of 50 CNA segments to the background signals with varied lengths (super short: 2~5 markers, short: 5~10 markers, medium: 10~20 markers, and long: 20~35 markers) and varied CNA proportions (20%, 40%, 60%, 80%, 100%), respectively. Deletion of a single copy (Del.s), mixed CNA states (mix) and duplication of a single copy (Dup.s) were spiked in separately. *F*1 score was utilized to evaluate the performance of CNA detection for each method.

**Supplementary Figure 8.**
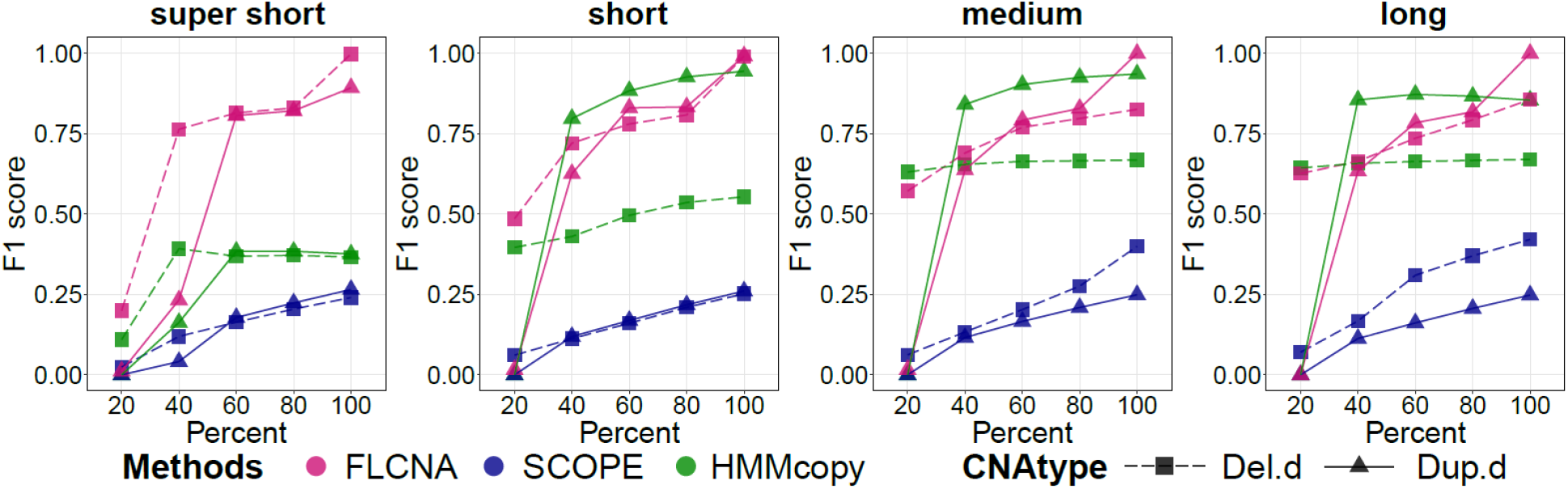
Assessment of FLCNA to detect CNAs using simulation data with three clusters and aberration of double copies. CNA calls were generated by FLCNA, SCOPE and HMMcopy, respectively. For each of three clusters, we added signals of 50 CNA segments to the background signals with varied lengths (super short: 2~5 markers, short: 5~10 markers, medium: 10~20 markers, and long: 20~35 markers) and varied CNA proportions (20%, 40%, 60%, 80%, 100%), respectively. Deletion of double copies (Del.d) and duplication of double copies (Dup.d) were spiked in separately. *F*1 score was utilized to evaluate the performance of CNA detection for each method.

**Supplementary Figure 9.**
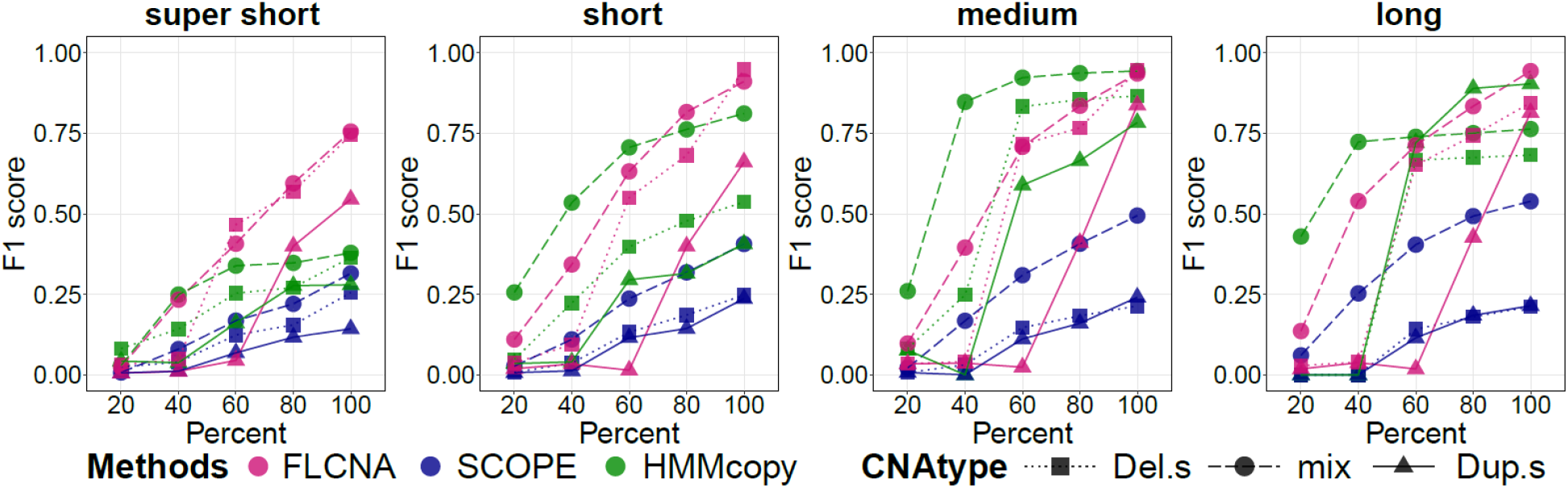
Assessment of FLCNA to detect CNAs using simulation data with five clusters and varied numbers of CNAs. CNA calls were generated by FLCNA, SCOPE and HMMcopy, respectively. For each of five clusters, we added signals of varied numbers of CNA segments (20~80) to the background signals with varied lengths (super short: 2~5 markers, short: 5~10 markers, medium: 10~20 markers, and long: 20~35 markers) and varied CNA proportions (20%, 40%, 60%, 80%, 100%), respectively. Deletion of a single copy (Del.s), mixed CNA states (mix) and duplication of a single copy (Dup.s) were spiked in separately. *F*1 score was utilized to evaluate the performance of CNA detection for each method.

**Supplementary Figure 10.**
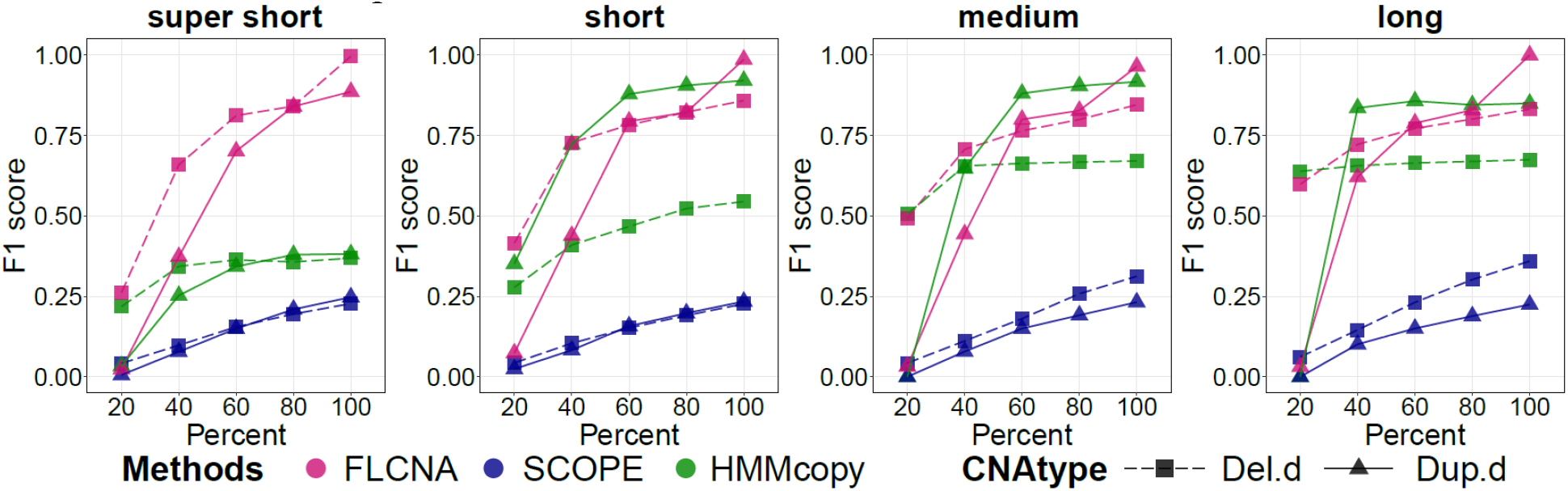
Assessment of FLCNA to detect CNAs using simulation data with five clusters, varied numbers of CNAs and aberration of double copies. CNA calls were generated by FLCNA, SCOPE and HMMcopy, respectively. For each of five clusters, we added signals of varied numbers of CNA segments (20~80) to the background signals with varied lengths (super short: 2~5 markers, short: 5~10 markers, medium: 10~20 markers, and long: 20~35 markers) and varied CNA proportions (20%, 40%, 60%, 80%, 100%), respectively. Deletion of double copies (Del.d) and duplication of double copies (Dup.d) were spiked in separately. *F*1 score was utilized to evaluate the performance of CNA detection for each method.

**Supplementary Figure 11.**
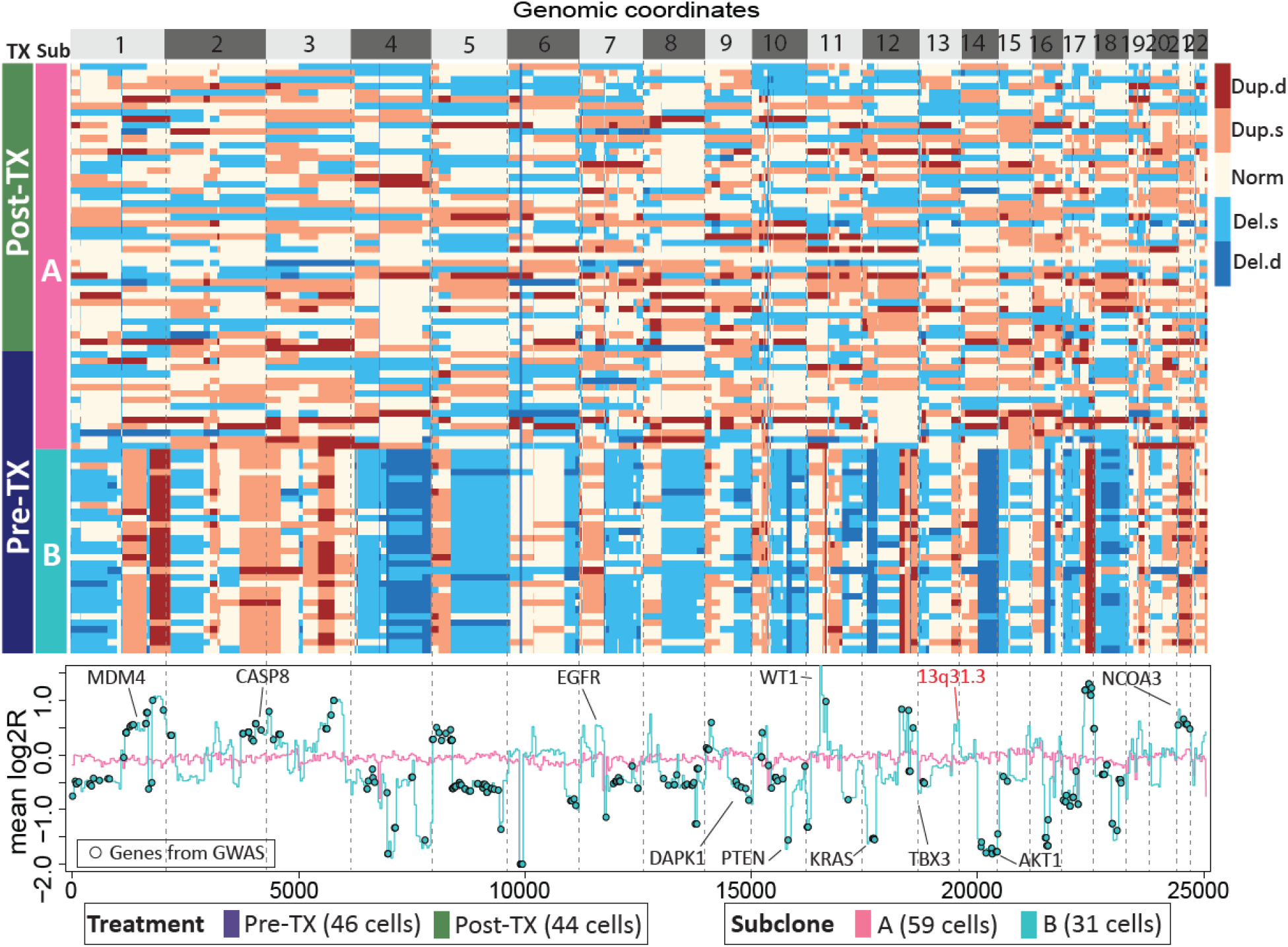
Subclone clustering of KTN129 patient using FLCNA. Cell clusters and copy number profile with different CNA states (Del.d, Del.s, Norm, Dup.s and Dup.d) were generated using FLCNA. Mean log2R were provided for each cluster. Shared CNAs identified using FLCNA were matched to significant genes from genome-wide association studies (GWAS) in the NHGRI-EBI GWAS Catalog. Del.d: Deletion of double copies; Del.s: Deletion of a single copy; Norm: Normal/diploid; Dup.s: Duplication of a single copy; Dup.d: Duplication of double copies; log2R: Logarithm transformation of ratio between normalized read counts and its sample specific mean; Pre-TX: pre-treatment; Post-TX: post-treatment.

**Supplementary Figure 12.**
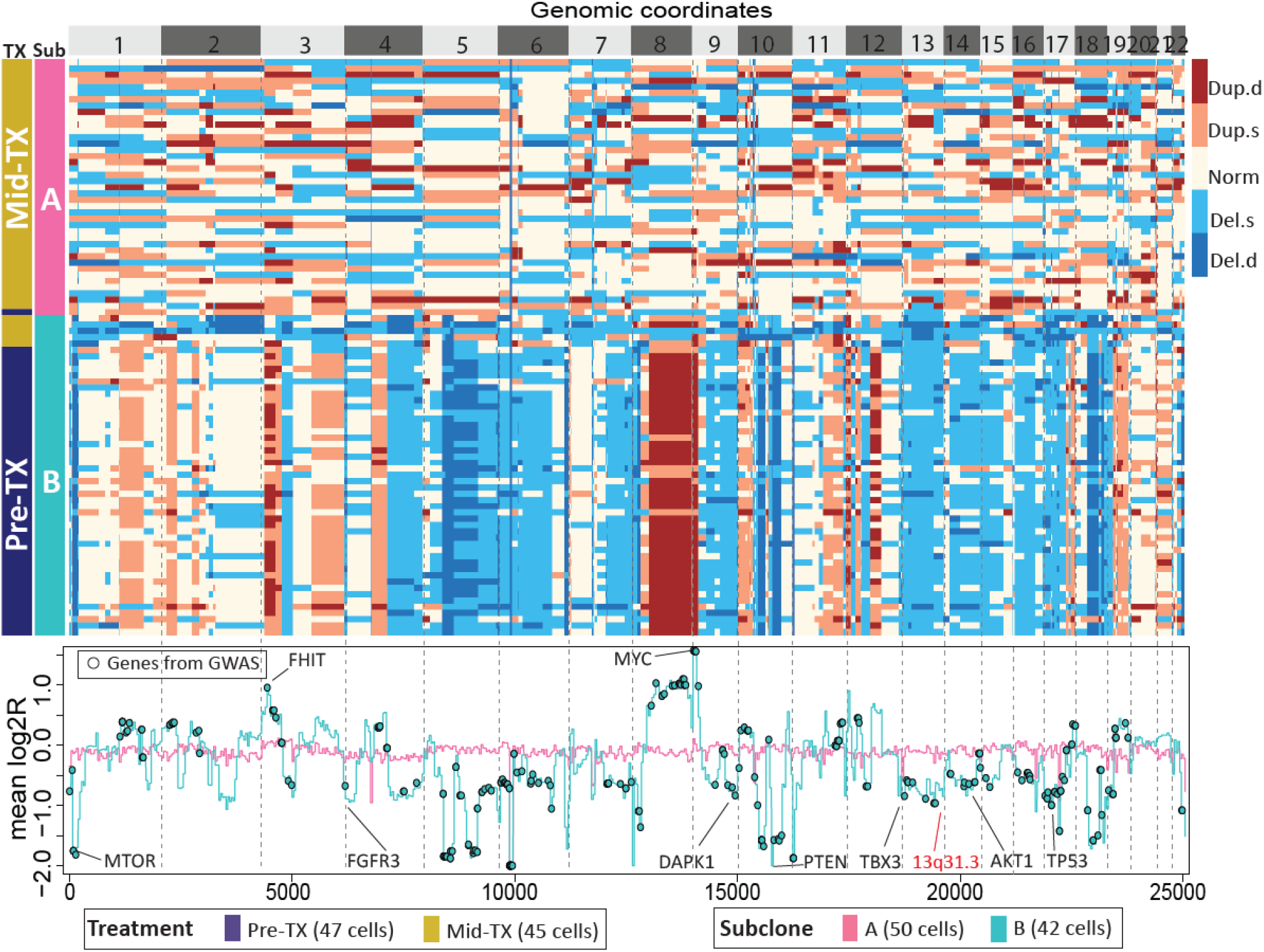
Subclone clustering of KTN302 patient using FLCNA. Cell clusters and copy number profile with different CNA states (Del.d, Del.s, Norm, Dup.s and Dup.d) were generated using FLCNA. Mean log2R were provided for each cluster. Shared CNAs identified using FLCNA were matched to significant genes from genome-wide association studies (GWAS) in the NHGRI-EBI GWAS Catalog. Del.d: Deletion of double copies; Del.s: Deletion of a single copy; Norm: Normal/diploid; Dup.s: Duplication of a single copy; Dup.d: Duplication of double copies; log2R: Logarithm transformation of ratio between normalized read counts and its sample specific mean; Pre-TX: pre-treatment; Mid-TX: mid-treatment.

**Supplementary Figure 13.**
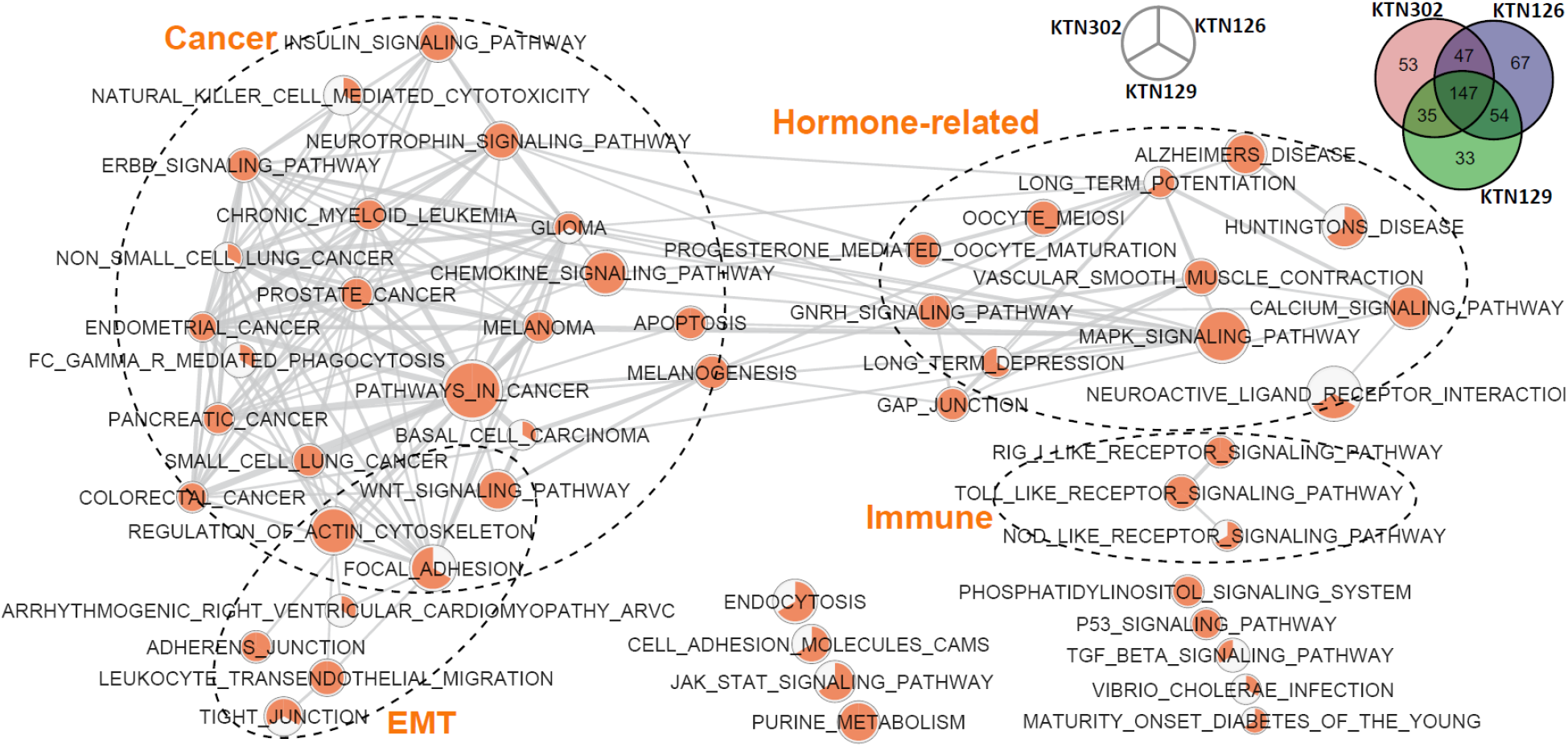
Gene expression networks in the TNBC dataset. The shared CNAs identified using FLCNA were mapped into significant genes from the genome-wide association studies (GWAS) with breast cancer. These matched genes were utilized for KEGG pathway enrichment analysis for three patients (i.e., KTN126, KTN129, KTN302). Each node in network is a pie plot showing three patients. Node size corresponds to the number of genes within the pathway. Colors inner the node correspond to the index whether this pathway is identified in this patient. Edge weight corresponds to the number of genes found in both connected pathways. Venn diagrams show the distribution of genes from GWAS which were also detected from above three patients. EMT: epithelial-mesenchymal transition.

**Supplementary Figure 14.**
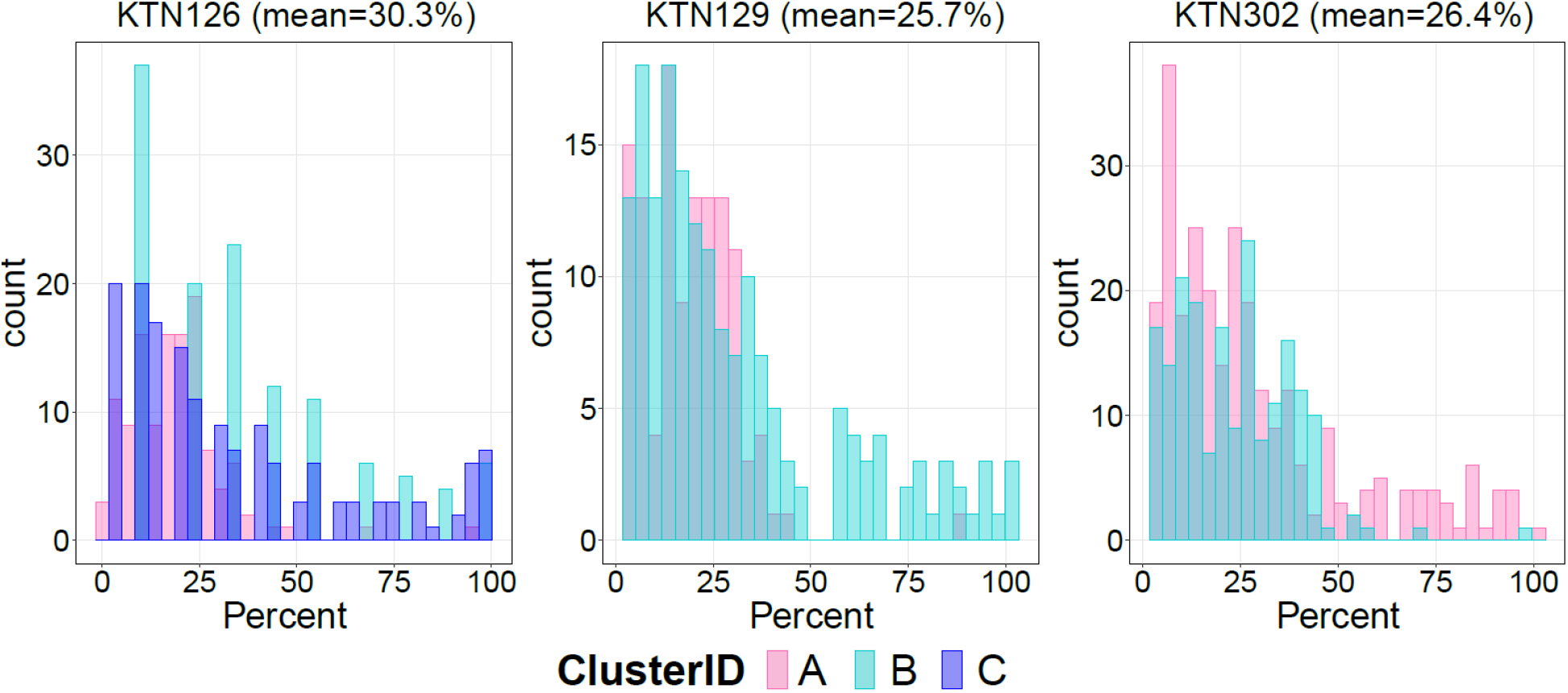
Distribution of shared percentage for CNAs detected using FLCNA in the TNBC dataset. CNAs were identified from the TNBC dataset with three patients (KTN126, KTN129, KTN302) using the FLCNA method.

**Supplementary Figure 15.**
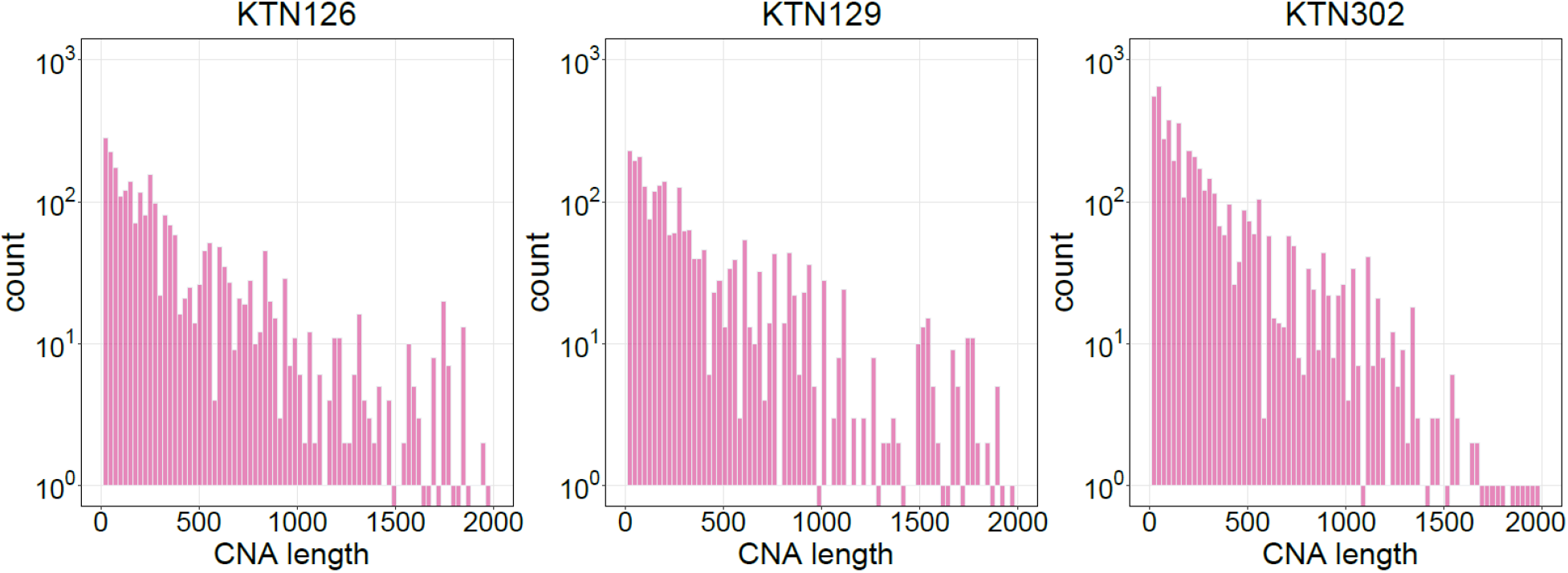
Distribution of CNAs detected using FLCNA in the TNBC dataset. CNAs were identified from the TNBC dataset with three patients (KTN126, KTN129, KTN302) using the FLCNA method.

## Supplementary Tables

**Supplementary Table 7.** Computational time of different CNA detection methods with scDNA-seq data. A high-performance cluster with 8 cores and 12GB RAM was used for CNA detection with KTN126 patient in the THBC dataset.

| Methods | Time (hours) |
| --- | --- |
| FLCNA | 1.20 |
| SCOPE | 10.5 |
| HMMcopy | 0.15 |

